# *Picornaviridae* and *Caliciviridae* diversity in Madagascar fruit bats is driven by cross-continental genetic exchange

**DOI:** 10.1101/2024.12.31.630946

**Authors:** Gwenddolen Kettenburg, Hafaliana C. Ranaivoson, Angelo Andrianianina, Santino Andry, Amy R. Henry, Rachel L. Davis, Farida Laboune, Elizabeth R. Longtine, Sucheta Godbole, Sophia Horigan, Emily Cornelius Ruhs, Vololoniaina Raharinosy, Tsiry Hasina Randriambolamanantsoa, Vincent Lacoste, Jean-Michel Heraud, Philippe Dussart, Daniel C. Douek, Cara E. Brook

**Affiliations:** Department of Ecology and Evolution, University of Chicago, IL, United States; Association Ekipa Fanihy, Antananarivo, Madagascar; Department of Zoology and Animal Biodiversity, University of Antananarivo, Antananarivo, Madagascar; Department of Entomology, University of Antananarivo, Antananarivo, Madagascar; Human Immunology Section, Vaccine Research Center, NIAID, NIH, Bethesda, USA; PREMISE, Vaccine Research Center, NIAID, NIH, Bethesda, USA; Grainger Center for Bioinformatics, Field Museum of Natural History, Chicago, IL, USA; Virology Unit, Institut Pasteur de Madagascar, Antananarivo, Madagascar

## Abstract

Bats are reservoir hosts for numerous well-known zoonotic viruses, but their broader virus-hosting capacities remain understudied. *Picornavirales* are an order of enteric viruses known to cause disease across a wide range of mammalian hosts, including Hepatitis A in humans and foot-and-mouth disease in ungulates. Host-switching and recombination drive the diversification of *Picornavirales* worldwide. Divergent *Caliciviridae* and *Picornaviridae* (families within the *Picornavirales*) have been described in bats across mainland Africa, but surveillance for these viruses has been rare in the Southwest Indian Ocean Islands. Bats live in close proximity to and are consumed widely as a food source by humans in Madagascar, providing opportunities for zoonotic transmission. Prior work in Madagascar has described numerous evolutionarily divergent bat viruses, some with zoonotic potential. Using metagenomic Next Generation Sequencing of urine and fecal samples obtained from three species of endemic Malagasy fruit bats (*Eidolon dupreanum*, *Pteropus rufus*, and *Rousettus madagascariensis*), we recovered 13 full-length and 37 partial-length genomic sequences within the order *Picornavirales* (36 *Picornaviridae* and 14 *Caliciviridae* sequences), which we identify and describe here. We find evidence that genetic exchange between mainland African bat and Madagascar bat *Picornavirales* likely shaped the diversification patterns of these novel sequences through recombination events between closely related *Picornavirales*; thus far, high host fidelity appears to have limited these viruses from spilling over into other species.

## INTRODUCTION

*Picornavirales* is a viral order associated with a large taxonomic variety of hosts, spanning animals (both vertebrates and invertebrates), plants and insects. Viruses in this order are characterized by a single-stranded, positive-sense RNA genome that forms non-enveloped icosahedral virions^1^. These viruses can encode either one or two polyproteins^1^. *Picornavirales* families *Picornaviridae* and *Caliciviridae* are known to cause clinical disease in both human and other mammalian hosts. Hepatitis A, caused by hepatovirus A in the *Picornaviridae* family, is a human disease with about 1.5 million reported cases each year, characterized by symptoms ranging from nausea and vomiting to potentially fatal fulminant hepatitis^2^. Poliovirus, an enterovirus in the family *Picornaviridae*, has caused epidemics of poliomyelitis, sometimes resulting in irreversible paralysis, throughout the 19^th^ and first half of the 20^th^ century before the introduction of the Salk and Sabin vaccines in the 1950s^3^. Although poliovirus disease met the target of 99% global eradication in 2000, poliovirus outbreaks still occur in areas of low vaccine uptake^4^. *Norovirus* and *Sapovirus*, in the family *Caliciviridae*, cause gastroenteritis in young children and more severe complications in immunocompromised individuals^5,6^.

Notably, viruses in the family *Picornaviridae* and *Caliciviridae* can infect some of the most common zoonotic hosts (bats, rodents, shrews)^7,8^, in addition to agriculturally significant hosts such as swine and cattle – both of which come into direct contact with humans^9–14^. Foot- and-mouth disease, caused by foot-and-mouth disease virus in the *Picornaviridae* family, affects cloven-hoofed animals, is highly contagious, and is characterized by clinical manifestations of vesicles in the oral cavity and feet^15^. Some *Picornaviridae* and *Caliciviridae* viruses display close genetic similarity between those hosted by humans and animals, but evidence of zoonoses is limited^14,16–19^. For example, some bat calicivirus virus-like particles (VLPs) experimentally generated *in vitro* have similar antigenic epitopes, which elicit histo-blood group binding, as do human and other mammalian noroviruses, but no natural observations of zoonoses for these viruses is known^14^. This diverse host range of *Picornaviridae* and *Caliciviridae* diversity can partially be equated to recombination and host-switching events that drive the evolution of these viruses^20–25^.

Bats have garnered interest for their unique ability to host viruses known to be highly pathogenic in other mammals, including humans, without experiencing significant disease^26–31^. While bats are known to host viruses in the *Picornavirales* order, particularly in the *Picornaviridae* and *Caliciviridae*^14,32–39^ families, these viruses are generally understudied as compared with a few more well-known bat virus clades such as coronaviruses, filoviruses, lyssaviruses and paramyxoviruses. Previous work from Cameroon used metagenomic Next Generation Sequencing (mNGS) to describe diverse *Picornavirales* in both *Eidolon helvum* and *Epomorphus gambianus* fruit bats, including novel kunsagiviruses and sapeloviruses that were divergent enough in amino acid composition to represent new species^36^. Divergent sapoviruses were also identified from the same bats^35^; in other cases, animal-derived sapoviruses have been shown to cluster closely with human-infecting genotypes^40–43^. In a separate study focusing on Algerian bats, evidence of past recombination events was detected within the genome of a novel mischivirus^34^. Host-virus coevolutionary analysis indicated that host-switching likely drove the diversification of this novel virus, in concordance with previously described patterns for the *Picornaviridae* family at large^34,44^. To date, the majority of bat *Picornavirales* work has taken place in mainland Africa, with limited prior studies in the Southwest Indian Ocean Islands (SWIO: Madagascar, Seychelles, Mauritius, Réunion, and Comoros)^13^.

Of the SWIO, Madagascar is particularly unique. An island country 400km off the coast of southeastern Africa, Madagascar boasts high levels of endemism and extraordinary evolutionary divergence among its flora and fauna due its isolation from mainland Africa and Asia for the past 80 million years^45^. Several evolutionarily distinct viruses have been previously identified in endemic Malagasy bats^46–52^, matching expectations that the isolated evolutionary landscape leads not only to diverse hosts but also diverse viruses. However, research regarding bat-hosted *Picornavirales* in Madagascar has been limited to only two prior studies. One of these studies described a unique hepatovirus in liver tissue collected from the Malagasy insectivorous bat, *Miniopterus cf. manavi*; sequence analysis of small mammal-hosted hepatoviruses broadly suggests some degree of host-virus co-evolution in their evolutionary history, with a hypothesized ancestral origin in bats and shrews^13^. In the other study, one full-length kobuvirus sequence was detected via mNGS of fecal samples collected from an *Eidolon dupreanum* fruit bat^52^. Within Madagascar, host-switching, likely fostered by co-roosting, is the dominant evolutionary mechanism driving diversification of morbilli-related bat paramyxoviruses^49^. We aimed to elucidate the role of host-switching mechanisms in driving the diversification of bat-borne *Picornaviridae* and *Caliciviridae* in Madagascar.

## MATERIALS AND METHODS

### Sample collection

Longitudinal monthly sampling of three endemic Malagasy fruit bats (*E. dupreanum, P. rufus,* and *Rousettus madagascariensis*) was carried out from 2013-2019 at species-specific roost sites across Madagascar in part with an ongoing effort to investigate seasonal viral dynamics, as described previously^46,47,51–53^. Over this period, 2156 bats were captured and processed under manual restraint as previously described^46,47,51,53^. Upon capture, bats were identified by species, age class, and sex, and individual fecal, throat and urine swabs were collected. All excreta samples were collected in viral transport medium (VTM) and frozen in liquid nitrogen until samples could be processed. A subset of 810 (271 fecal/539 urine) samples collected between 2013-2019 at the following roost sites was used for the molecular analyses outlined here: Angavobe cave (-18.944S, 47.949 E, *E. dupreanum)*; Angavokely cave (-18.933 S, 47.758 E, *E. dupreanum)*; Ambakoana roost (-18.513 S, 48.167 E, *P. rufus)*; Maromizaha cave (-18.9623 S, 48.4525 E, *R. madagascariensis)*; Andrafiabe cave (-12.9435 S, 49.0555 E, *E. dupreanum* and *R. madagascariensis*); Cathedral cave (-12.952901 S, 49.046885 E, *E. dupreanum*); Antsiroandoha cave (-12.959336 S, 49.123698 E, *E. dupreanum*). This study was carried out in strict accordance with research permits obtained from the Madagascar Ministry of Forest and the Environment (permit numbers: 251/13, 166/14, 75/15, 92/16, 259/16, 019/18, 170/18, and 007/19,) and under guidelines posted by the American Veterinary Medical Association. All field protocols employed were pre-approved by the UC Berkeley Animal Care and Use Committee (IACUC Protocol # AUP-2017-10-10393), and every effort was made to minimize discomfort to animals.

### Sample processing

Samples were frozen at the site of capture in liquid nitrogen and stored at -80°C at the Virology Unit at the Institut Pasteur de Madagascar. Subsequently, RNA extraction of the samples was completed using the Zymo Quick DNA/RNA Microprep Plus kit (Zymo Research, Irvine, CA, USA), according to the manufacturer’s instructions and including the step for DNAse digestion. Extracted RNA was then shipped to either the Chan Zuckerberg Biohub (CZB; sample date range 2018-2019) (San Francisco, CA, USA) or the Vaccine Research Center (VRC), National Institute of Allergy and Infectious Diseases (NIAID), National Institutes of Health (NIH; sample date range 2013-2019) (Bethesda, USA) for mNGS.

Briefly, RNA underwent library preparation using the NEBNext Ultra II RNA Library Prep Kit (New England Biolabs, Beverly, MA, USA) with the following modifications for CZB samples (date range 2018-2019): 25pg of External RNA Controls Consortium Spike-in mix (ERCCS, Thermo-Fisher) was added to each sample prior to RNA fragmentation; the input RNA mixture was fragmented for 8min at 94°C prior to reverse transcription; and a total of 14 cycles of PCR with dual-indexed TruSeq adapters was applied to amplify the resulting individual libraries. Quality was assessed by electrophoresis before performing large-scale paired-end sequencing (2 x 146bp) on the Illumina NovaSeq (Illumina, San Diego, CA, USA). The following modifications were made for VRC samples (date range 2018-2019): input RNA was fragmented for 7min at 94°C prior to reverse transcription; and a total of 12 cycles of PCR. Quality was assessed by electrophoresis (Bioanalyzer 2100, Agilent) before paired-end sequencing (2 x 150bp) on the Illumina NovaSeq 6000 (Illumina, San Diego, CA, USA).

The CZID pipeline to parse output from individual libraries into FASTQ format is available on GitHub (https://github.com/czbiohub/utilities).

### Virus detection

Raw reads were host-filtered, quality-filtered, and assembled on the Chan Zuckerberg Infectious Diseases (CZID) bioinformatics platform (v3.10 NR/NT 2019-12-01)^54^. A background profile named “bat” was created using all publicly available full-length bat genomes in GenBank at the time of sequencing (July 2019 for CZID samples, December 2023 for VRC samples) and used in host filtering. Samples were investigated using a command line BLAST pipeline, as described previously in novel coronavirus, henipavirus, kobuvirus, and astrovirus detection^46,47,51,52^. The following criteria were used to call “positive” samples: (1) at least two contigs with an average read depth of >2 reads/nucleotide were assembled and (2) showed significant nucleotide or protein BLAST alignment(s) (alignment length >100nt/aa and E-value < 0.00001 for nucleotide BLAST/bit score >100 for protein BLAST) to *Picornavirales* present in NCBI NR/NT database (v12-01-2019). We used NCBI Virus taxids 12058/478825 for *Picornaviridae*/unclassified *Picornaviridae* and taxids 11974/179239 for *Caliciviridae*/unclassified *Caliciviridae* to develop nucleotide and protein libraries, against which we queried our novel sequences in command line BLASTn^55^ and BLASTx^55^ searches. Detailed instructions on our methods for parsing “positive” samples is available on our open-access GitHub repository (see **Data Availability**).

### Genome annotation

Once contigs were identified as *Picornaviridae*/*Caliciviridae* sequences, we sorted genomes into full and partial-length sequences. We downloaded annotated reference *Picornaviridae*/*Caliciviridae* sequences from NCBI and aligned to our novel sequences using MAFFT^56^ (v7.450) with default settings in Geneious Prime (v08-18-2022). We annotated the polyprotein and peptide sequences using available *Picornaviridae*/*Caliciviridae* reference sequences that corresponded to the top BLAST hit. Reference sequences were also used to identify potential cleavage sites and conserved motifs within RNA-dependent RNA polymerase (RdRp), polyprotein major proteases, and NTPase-helicase.

### Phylogenetic analysis

We constructed ten maximum-likelihood (ML) nucleotide phylogenetic trees in RAxML^57^, representing all genera in which we identified at least one novel sequence >2000bp in length. All trees were rooted using an outgroup of Sindbis virus (accession NC_001547). We first constructed (A) a phylogeny encompassing the conserved polymerase (3D peptide) region and included global background sequences from viral genera corresponding to those clades represented by all novel Madagascar sequences (unclassified bat picornavirus, *Cardiovirus*, *Hepatovirus*, *Kobuvirus*, *Kunsagivirus*, *Mischivirus*, *Sapelovirus*, *Sapovirus*, and *Teschovirus*). Then, we constructed nine additional ML trees using full-length sequences from the following viral genera as references: (B) *Cardiovirus*, (C) *Hepatovirus*, (D) *Kobuvirus*, (E) *Kunsagivirus*, (F) *Mischivirus*, (G) *Shanbavirus*/unclassified bat picornavirus, (H) *Sapelovirus*, (I) *Teschovirus*, and (J) *Sapovirus*. Reference sequences and novel sequences were aligned using MAFFT^56^, and we used ModelTest-NG^58^ (v0.1.7) to determine the best nucleotide substitution model for each alignment prior to building the ML-trees. We further detail NCBI virus taxid information, ModelTest output, alignment names, and size of overlap region per phylogeny in **Supplemental table 1**. Sequence alignments and more detailed tree building instructions are available in our GitHub repository (see **Data Availability**).

### Similarity analysis

We conducted similarity analyses, using PySimPlot^59^, on all nucleotide and translated amino acid *Picornaviridae* and *Caliciviridae* full-length sequences recovered from CZID. PySimPlot commands were formulated using our novel Madagascar sequences as the query sequences; in cases where multiple novel sequences were identified within a single genus, we input one of those sequences as the query and included the others within the reference sequences. Otherwise, reference sequences for input to PySimPlot^59^ were comprised of the top three full-genome BLASTx hits for the query. In all cases, sequences were aligned in MAFFT^56^, and both similarity analyses were carried out using a window size of 100aa and a step size of 20aa for amino acid comparisons and a window size of 100bp and a step size of 1bp for nucleotide comparisons. A full list of query sequences and reference sequences per alignment is available in our GitHub repository (see **Data Availability**).

### RDP4 recombination analysis

Recombination Detection Program 4 (RDP4)^60^ was used to evaluate signals of recombination in sets of pairwise alignments; RDP4 identifies the potential recombinant sequence, as well as the potential major parental sequence (from which most of the genome is reputed to be sourced) and the minor parental sequence (from which some of the genome is reputed to be sourced) for this lineage. From each pairwise alignment, RDP4 identifies potentially recombinant sequences, in addition to specifying the region of each putatively recombinant genome most likely to have undergone recombination and the parent genome (either major or minor) most likely to have contributed to that region. We ran seven separate analyses within RDP4 (RDP, GENECONV, Bootscan, MaxChi, Chimaera, and 3Seq) to estimate likely recombination events and the location of the recombination event within all full-length (13) sequences identified in this study. We considered recombination events to be reliable if at least four analyses were significant using default settings with a cutoff *P*-value of 0.05. A full list of query sequences and reference sequences per alignment is available in our GitHub repository (see **Data Availability**).

## RESULTS

### Sampling and demographic patterns

We sequenced RNA extracted from 810 (271 fecal and 539 urine) samples from 803 individual bats (373 *E. dupreanum*, 146 *P. rufus*, and 284 *R. madagascariensis*). In total, 24/803 bats were *Picornaviridae* positive (2.98%), and 7/803 bats were *Caliciviridae* positive (0.87%) (**Table 1**). Of the 24 *Picornaviridae* positive bats, 16 were *E. dupreanum*, 1 was *P. rufus*, and 7 were *R. madagascariensis*. The 7 *Caliciviridae* positive bats were evenly split between *E. dupreanum* (4) and *R. madagascariensis* (4) with no infections identified in *P. rufus*. No bats were positive at the Ankarana caves (N=151). *Picornaviridae* and *Caliciviridae* prevalence was highest in *E. dupreanum* at the Angavokely/Angavobe roosts (16/281 bats *Picornaviridae* and 4/281 bats *Caliciviridae*) with a similar prevalence reported in *R. madagascariensis* at the Maromizaha roost (7/225 bats *Picornaviridae* and 4/225 bats *Caliciviridae*) (**Fig. 1A and Table 1**). In *P. rufus*, only one bat out of 146 individuals was found to be positive for *Picornaviridae* (**Fig. 1A and Table 1**). Two individual bats were coinfected: one was an *E. dupreanum* from Angavokely/Angavobe cave, from which we identified a partial genome for a *Hepatovirus* and a full genome for the only *Kunsagivirus* described in this study. The second individual was a *R. madagascariensis* from Maromizaha, which hosted a partial genome for a *Sapovirus* and a full genome for an unclassified bat picornavirus.

**Figure 1:**
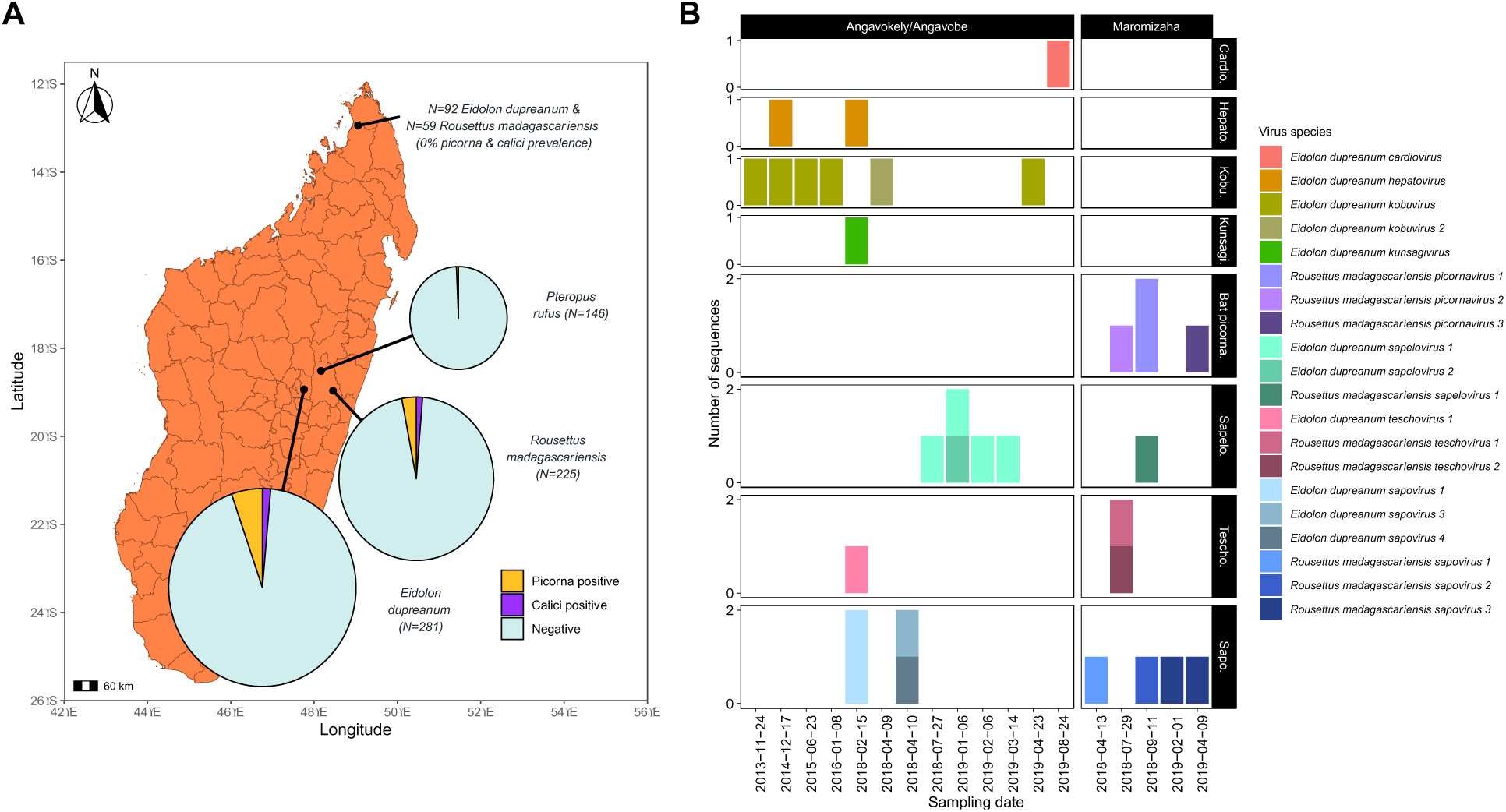
(A) Map of sampling sites for *Eidolon dupreanum, Pteropus rufus*, and *Rousettus madagascariensis* at sites in Districts of Ambilobe (*E. dupreanum* and *R. madagascariensis*: Ankarana caves), Moramanga (*P. rufus:* Ambakoana roost and *R. madagascariensis*: Maromizaha cave), and Manjakandriana (*E. dupreanum*: Angavobe/Angavokely caves), Madagascar. Pie charts show *Picornaviridae* (yellow) and *Caliciviridae* (orange) positive bats by site. Pie chart corresponds to sample size on a log10 scale. (B) Summary bar plot of diversity of viral genera (facets) and viral species (colors) identified from bats caught within the same sampling date at Angavokely/Angavobe caves (*E. dupreanum*) and Maromizaha cave (*R. madagascariensis*). Ambakoana roost (P*. rufus*) was excluded due to only having one identified novel virus in a single bat.

**Table 1:** Sampling efforts by roost site and species, in addition to breakdown by *Picornaviridae* and *Caliciviridae* positives.

| District | Roost Site | Species | # Tested Bats | *Picornaviridae* positive bats | *Caliciviridae* positive bats |
| --- | --- | --- | --- | --- | --- |
| Ambilobe, Madagascar | Ankarana caves (Andrafiabe, Cathedral, Antisiroandoha) | *E. dupreanum* | 92 | 0 | 0 |
|  |  | *R. madagascariensis* | 59 | 0 | 0 |
| Moramanga, Madagascar | Ambakoana roost | *P. rufus* | 146 | 1 (0.68%) | 0 |
|  | Maromizaha cave | *R. madagascariensis* | 225 | 7 (3.11%) | 4 (1.77%) |
| Manjakandriana, Madagascar | Angavobe/Angavokely caves | *E. dupreanum* | 281 | 16 (5.69%) | 4 (1.42%) |

### Genome characterization and roost-site diversity

We recovered 13 full-length (12 *Picornaviridae* and 1 *Caliciviridae*) and 37 partial-length (24 *Picornaviridae* and 13 *Caliciviridae*) sequences primarily from *E. dupreanum* (25 *Picornaviridae* from 15 individuals and 5 *Caliciviridae* from 4 individuals) and *R. madagascariensis* (10 *Picornaviridae* from 7 individuals and 9 *Caliciviridae* from 4 individuals) (**Fig. 1B**). Only one viral *Picornaviridae* sequence (full-genome) was recovered from an individual *P. rufus* (**Fig. 1B**).

Within *Picornaviridae*, we identified sequences corresponding to the following viral genera: *Cardiovirus* (1 sequence from 1 viral species), *Hepatovirus* (4 sequences from 1 viral species), *Kobuvirus* (8 sequences from 2 viral species), *Kunsagivirus* (1 sequence from 1 viral species), *Mischivirus* (1 sequence from 1 viral species), *Sapelovirus* (11 sequences from 3 viral species, *Teschovirus* (3 sequences from 3 viral species), and unclassified “bat picornavirus” (7 sequences from 3 viral species). *Caliciviridae* was only represented by a single genus: *Sapovirus* (14 sequences from 7 viral species) (**Fig. 1B, Supplemental tables 2** and **3**).

Some *Picornaviridae* genera were only found in *E. dupreanum* (*Cardiovirus*, *Hepatovirus*, *Kobuvirus*, and *Kunsagivirus*) (**Fig. 1B**, **Supplemental table 2**). In general, *Picornaviridae* sequences from *E. dupreanum* had highest identity to *Picornaviridae* identified from its sister species *E. helvum* (**Supplemental table 2**), which is widely distributed across the African mainland continent but is absent from Madagascar^61^. Exceptions to this pattern included the *Cardiovirus* sequence which had the highest identity to a divergent encephalomyocarditis virus isolated from an orangutan from Singapore (NCBI accession QMI58083.1)^62^ and a *Teschovirus* sequence with highest identity to a Ugandan *Rousettus aegypticus bat teschovirus* (accession XBH24020)^63^. *Cardiovirus* has been detected in bats before, namely East Asian *Miniopterus fuliginosus* bats^64^. It is possible that our novel Malagasy bat *Cardiovirus* would be more closely related to this *Chiropteran*-hosted *Cardiovirus;* however, the lack of publicly-available sequences from this previous study impedes these comparisons. One partial *Teschovirus* sequence from a Saudi Arabian *E. helvum* bat has also been described (accession KX420938)^65^, but is still not the highest identity to our novel *E. dupreanum*-derived *Teschovirus*. *E. dupreanum*-hosted *Hepatovirus*, *Kobuvirus*, *Kunsagivirus,* and *Sapelovirus* had highest identity to Ghanaian *E. helvum*-hosted hepatovirus H2 (accession YP_009179216.1)^13^, Malagasy *E. dupreanum*-hosted kobuvirus (accession WBP49885.1)^52^, Cameroonian *E. helvum*-hosted kunsagivirus B (accession YP_009345896.1)^66^, and Cameroonian *E. helvum*-hosted sapelovirus (accession YP_009345901.1)^66^ respectively (**Supplemental table 2**).

One *Picornaviridae* genus, *Mischivirus*, was found only in *P. rufus,* in one individual (**Fig. 1B**). This novel sequence showed very low identity (46% over 81% genome coverage) to its closest match (accession YP_009121743.1, mischivirus C1 from a *Hipposideros gigas* bat from the Democratic Republic of the Congo (DRC)) (**Supplemental table 2**).

Among *Picornaviridae* genera exclusively hosted by *R. madagascariensis* (unclassified bat picornavirus), and those hosted by both *E. dupreanum* and *R. madagascariensis* (*Sapelovirus* and *Teschovirus*), we identified numerous viruses with high identity to viruses hosted by sister species *R. aegypticus*^67^ sampled in Uganda and Kenya. The unclassified bat picornaviruses were most closely related to other previously-described “bat picornaviruses”^12,63,68,69^ but, within this clade, form what appears to be a largely divergent *Picornaviridae* genus with an average BLASTx identity of ∼80% with ∼90% genome coverage to the closest match (**Supplemental table 2**). *R. madagascariensis*-hosted *Sapelovirus* and *Teschovirus* had highest identity to East African (Uganda and Kenya) *R. aegypticus*-hosted *Sapelovirus* and *Teschovirus* sequences^63^ (**Supplemental table 2**).

The representative *Caliciviridae* genus, *Sapovirus*, was found in both *E. dupreanum* and *R. madagascariensis* (**Fig. 1**, **Supplemental table 3**). *Sapovirus* sequences from *E. dupreanum* showed highest identity to a Cameroonian *E. helvum*-hosted *Sapovirus* (accession KX759623.1)^35^ (**Supplemental table 3**) and *R. madagascariensis*-hosted *Sapovirus* sequences had highest identity to East African *R. aegypticus*-hosted sapoviruses, mirroring the host patterns seen in the novel *Picornaviridae* sequences, as well (**Supplemental table 3**). We identified as many as 4 different viral species sourced from the same group of bats captured within a single sampling session (**Fig. 1B**). Overall, the Angavokely roost site (with *E. dupreanum* bats) was represented the most frequently in the samples analyzed for this study and demonstrated the most unique viruses overall (**Fig. 1B** and **Supplemental tables 2** and **3**), but the Maromizaha roost site (with *R. madagascariensis* bats) also demonstrated high virus diversity – with multiple repeats of the same virus genotype recovered from different individuals – within fewer viral genera.

### Genome annotation

#### Picornaviridae

We successfully annotated a single ORF-encoding polyprotein in *Picornaviridae* which includes the P1 region of structural polypeptides and the P2/P3 regions of replication-associated nonstructural polypeptides) and further identified cleavage sites within the polyprotein between each peptide of the P1 region (L, VP4, VP2, VP3, VP1), the P2 region (2A, 2B, 2C) and the P3 region (3A, 3B, 3C, and the RNA-dependent-RNA-polymerase [RdRp] 3D) (**Supplemental table 4**). Following previous work^33,36,68^, we identified the conserved motifs, helicase GxxGxGKS, 2A protease GxCG, 3C protease GxCG, and RdRp motifs KDELR, YGDD, and FLKR. All novel *Picornaviridae* identified in our study had conserved RdRp motifs with no substitutions (**Table 2**). The only genera with the 2A protease motif were the *R. madagascariensis picornaviruses* and all sapeloviruses. *E. dupreanum cardiovirus* had the same 2C GDAGQGKS helicase as the previously mentioned divergent *simian encephalomyocarditis virus*^62^ (**Table 2**). *E. dupreanum* hepatoviruses all had the same 2C motif GNRGGGKS as the comparable Cameroonian *E. helvum hepatovirus* H2^13^ and a treeshrew hepatovirus (accession NC_028981) (**Table 2**). *E. dupreanum kobuvirus* sequences were generally almost identical to a previously described *E. dupreanum kobuvirus*^52^ with slight genotypic variation; the 2C helicase GPPGTGKS and 3C protease GLCG motifs were perfectly identical. We characterized a partial genome of a second kobuvirus species, *E. dupreanum kobuvirus* 2, but the segment unfortunately did not cover any of the conserved regions for further comparison. These motifs seem to be well conserved in kobuviruses across multiple host types such as bovine, human, canine, and other bats^70–73^ (**Table 2**). *E. dupreanum kunsagivirus* has the same 2C helicase GEPGTGKS and 3C protease GMCG as other African bat and rodent kunsagiviruses^66,74^.

**Table 2:**
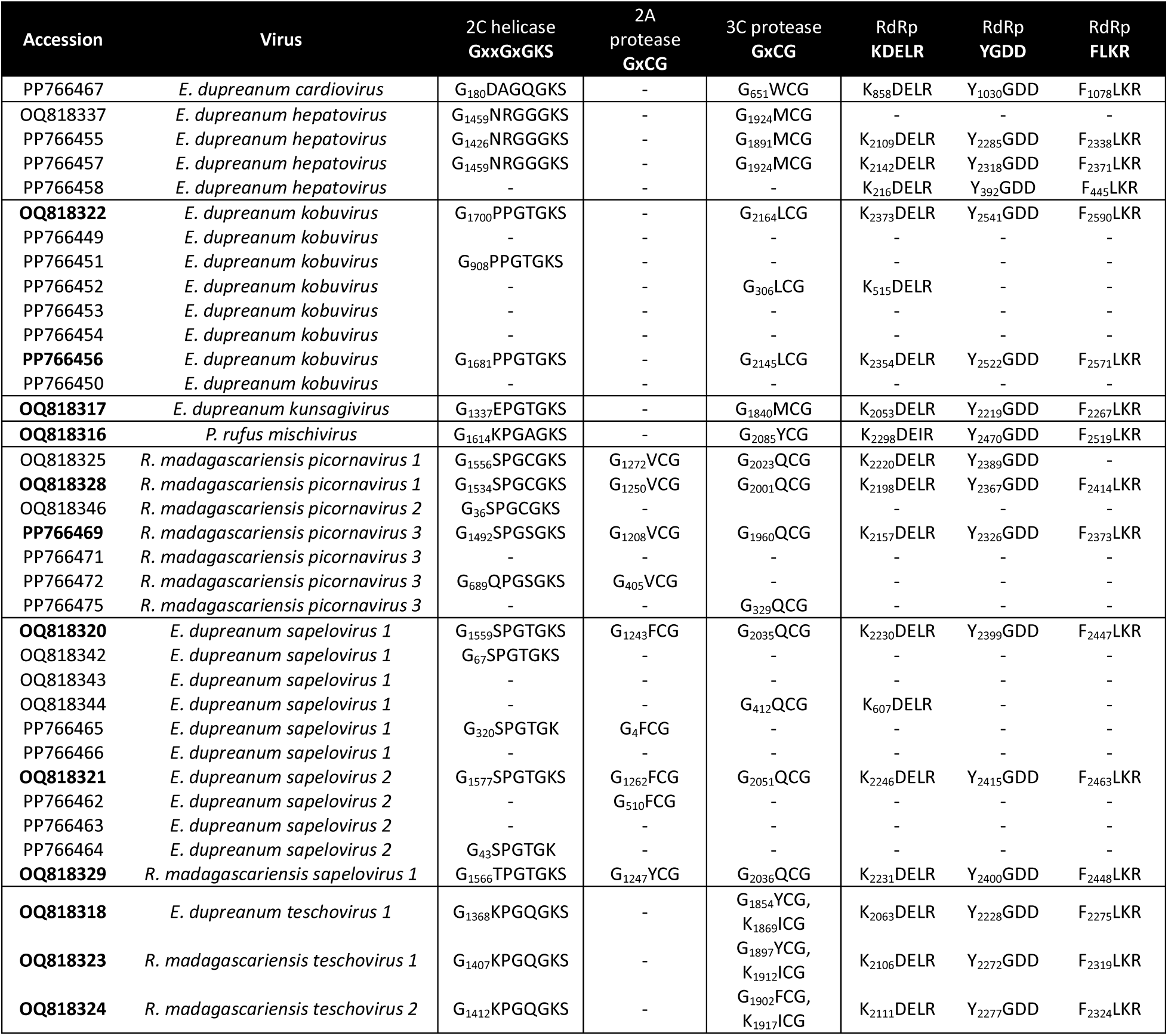
Conserved motifs in novel *Picornaviridae* sequences from Madagascar fruit bats. Bold denotes full-length sequences. Dashes indicate that the sequence recovered does not include that motif due to length. Absent motifs are otherwise noted.

The divergent *P. rufus mischivirus* had a 2C helicase motif GKPGAGKS and 3C protease motif GYCG as seen in Chinese *Miniopterus pusillus*-hosted mischiviruses (accession OR867092), and an Algerian *Miniopterus* spp.-hosted mischivirus (accession MG888045). While the closest match for *P. rufus* mischivirus is a mischivirus C1 sequence from *H. gigas* bat from the DRC, it did not share the 2C helicase of GRPGAGKS or the 3C protease of GFCG (**Table 2**.

Also divergent, the group of *R. madagascariensis*-hosted bat picornaviruses share the same 3C helicase motif GSPGCGKS, excepting one partial sequence of *R. madagascariensis picornavirus* 3 (accession PP766472) which instead had GQPGSGKS, a motif previously seen in *bat picornavirus* BtSY4 sequences from Chinese *Rhinolophus* bats (accessions PP746000 and OP963617)^69^. Otherwise, the GSPGCGKS motif is shared with East African *E. helvum* and *R. aegypticus picornaviruses* (sample accessions PP711943 and PP711928)^63^, as is the 2A protease GVCG motif (**Table 2**). Teschoviruses from *E. dupreanum* and *R. madagascariensis* had the 2C helicase motif of GKPGQGKS, as seen in Chinese *R. leschenaultii* bats (accessions OR951333 and OR951334), differing from the 2C helicase GAPGQGKS seen in East African *R. aegypticus* bats (accession PP711934) and porcine teschoviruses (sample accession OM105029). Teschoviruses lack a 2A protease motif but have two 3C protease motifs that are generally conserved across bat and porcine hosts (GYCG and KICG), excepting *R. madagascariensis teschovirus* 2, which had a novel substitution of GFCG in the first 3C protease motif (**Table 2**). Finally, the *E. dupreanum sapeloviruses* had a 2C helicase motif of GSPGTGKS and a 2A protease motif of GFCG, whereas *R. madagascariensis sapelovirus* 1 has a 2C helicase motif of GTPGTGKS and a 2A protease motif of GYCG (**Table 2**). East African *E. helvum*-hosted sapeloviruses have the same motifs as *E. dupreanum* sapeloviruses (accessions PP711921 and PP711943)^63^, and East African *R. aegypticus*-hosted sapeloviruses have the same motifs as *R. madagascariensis sapelovirus* 1 (accession PP711911)^63^, indicating bat species-specific viral differences that are conserved between sister species.

#### Caliciviridae

In all *Caliciviridae* sequences we identified and annotated the ORF1-encoding polyprotein in addition to ORF2. Within ORF1 polyprotein we further identified cleavage sites between peptides NS1/NS2, Helicase, NS4, Vpg, Pro-Pol (RdRp), and VP1 in addition to annotating the small structural protein VP2 encoded by ORF2 (**Supplemental table 5**).

Within *Caliciviridae*, specifically sapoviruses, again following prior work^35^, we identified conserved helicase GAPGIGKT, Vpg KGKTK and DDEYDE, protease GxCG, RdRp conserved WKGL, KDELR, DYSKWDST, GLPSG, and YGDD, and finally VP1 PPG and GWS motifs. RdRp and VP1 motifs were generally conserved among *E. dupreanum* sapoviruses and *R. madagascariensis* sapoviruses, in addition to all comparison sequences (**Table 3**). Slight variation existed among the Malagasy bat sapoviruses, notably a missing VpG KGKTK motif and a protease GSCG (other novel sequences are GDCG) in *R. madagascariensis sapovirus* 3: accession OQ818348 (**Table 3**). *R. madagascariensis sapovirus* 2: accession OQ818347 had a RdRp motif of DFSKWDST (other novel sequences have DYSKWDST), also seen in East African *R. aegypticus* sapoviruses (accessions PP712001 and PP712004). NTPase GPPGIGKT was well conserved amongst the novel sapoviruses, also appearing in African bat sapoviruses (example accessions KX759619 and PP712001), but also in a human sapovirus (accession MH922772) and a pig sapovirus (accession OM105025) (**Table 3**).

**Table 3:**
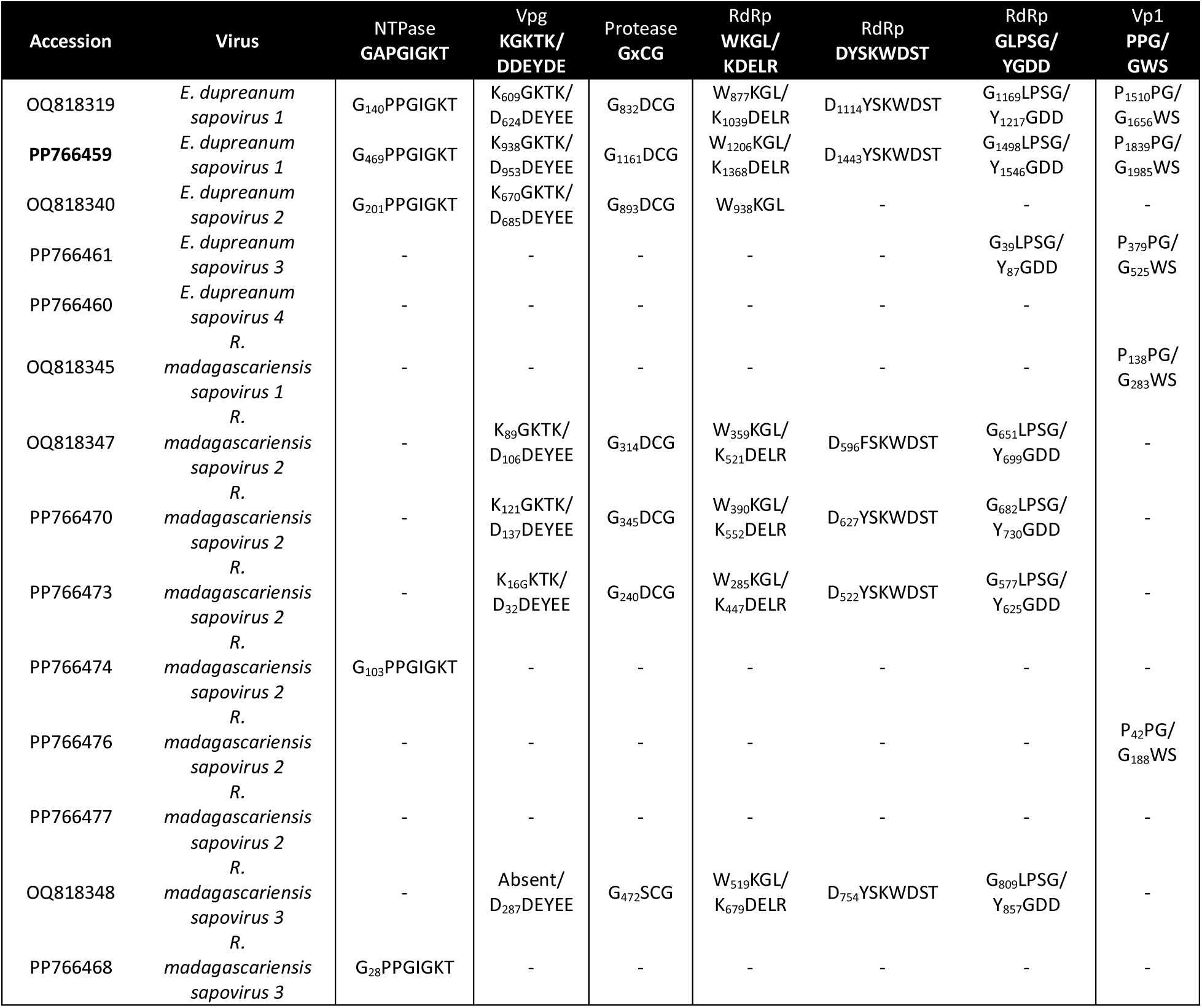
Conserved motifs in novel *Caliciviridae* sequences from Madagascar fruit bats. Bold denotes full-length sequences. Dashes indicate that the sequence recovered does not include that motif due to length. Absent motifs are otherwise noted.

### Phylogenetic analysis

#### Summary phylogeny of Picornaviridae and Caliciviridae

After identification of viral genera and species through BLAST (**Supplemental tables 2** and **3**), we constructed an RdRp phylogeny which showed phylogenetic clustering of the novel Malagasy bat *Picornaviridae* and *Caliciviridae* with other bat-hosted sequences (**Fig. 2**). For genomes recovered, read support was sufficient (average support >88 reads/base pair in full genomes, average support >1764 reads/base pair in partial genomes) (**Supplemental Fig. 1** and **2**). In all cases, support was lower in the 5’ and 3’ untranslated regions (UTRs) (**Supplemental Fig. 1** and **2**). *Cardiovirus* (average support 12007 reads/base pair) recovered the most read support (although, only in a small portion of the genome) followed by *Mischivirus* (average support 712 reads/base pair), *Sapovirus* (average support 635 reads/base pair) and *Kobuvirus* (average support 174 reads/base pair) (**Fig. 2, Supplemental Fig. 1** and **2**). While lower, read support was sufficient for *Teschovirus* (average support 70 reads/base pair), *Kunsagivirus* (average support 29 reads/base pair), *Sapelovirus* (average support 16 reads/base pair), *Hepatovirus* (average support 15 reads/base pair), and bat picornaviruses (average support 13 reads/base pair) (**Fig. 2, Supplemental Fig. 1** and **2**). We further analyzed assembled contigs (contiguous sequences representing partial or full genomes submitted to NCBI) in their own genus-specific phylogenies (**Fig. 3A-I**). We recovered the most contigs from the genus *Sapovirus*, followed by *Sapelovirus* and bat picornaviruses (**Fig. 2**). Though read support was high for *Kobuvirus* and *Cardiovirus* sequences identified, we found fewer discrete contigs representing unique virus species from different individuals for these taxa (**Fig. 2**).

**Figure 2:**
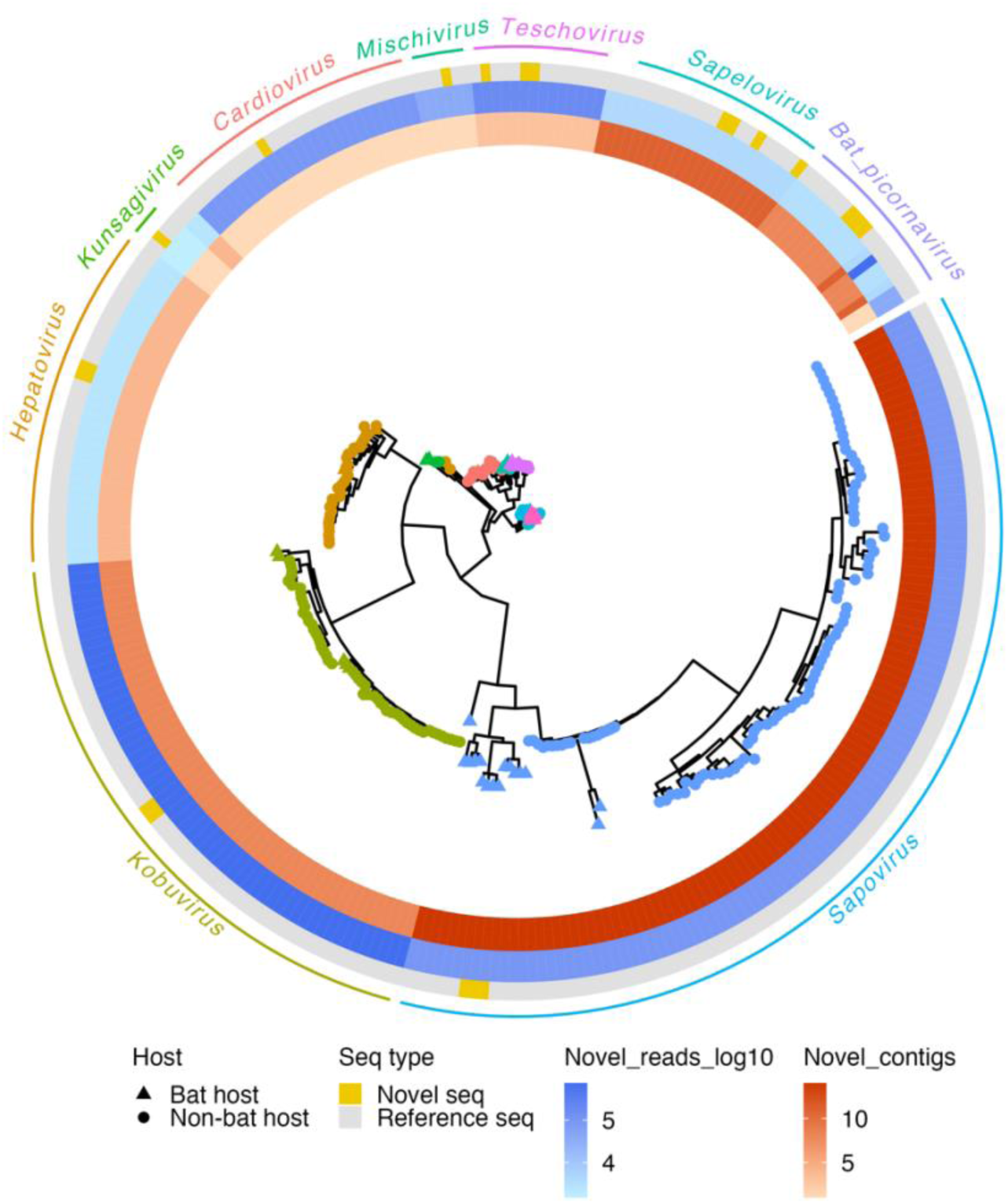
Maximum-likelihood nucleotide phylogeny (RAxML-NG-MPI^57^) of sequences from *Picornaviridae* and *Caliciviridae* genera from which novel sequences were classified from a 2900 bp overlapping region of the RdRp region of the genome, using a best-fit GTR+I+G4 nucleotide substitution model (**Supplemental table 1**). Bootstraps were computed using Felsenstein’s method and are visualized on tree branches. Rings indicate number of contigs (red) and reads (blue and log10 scale) recovered for each novel sequence genera. Novel sequences are indicated by a yellow square. Tip shape indicates host in which the sequence was derived corresponding to the legend. Tree is rooted in Sindbis virus (accession NC_001547.1). Roots have been removed for ease of visualization. Branch lengths are scaled by nucleotide substitutions per site, corresponding to scalebar.

**Figure 3:**
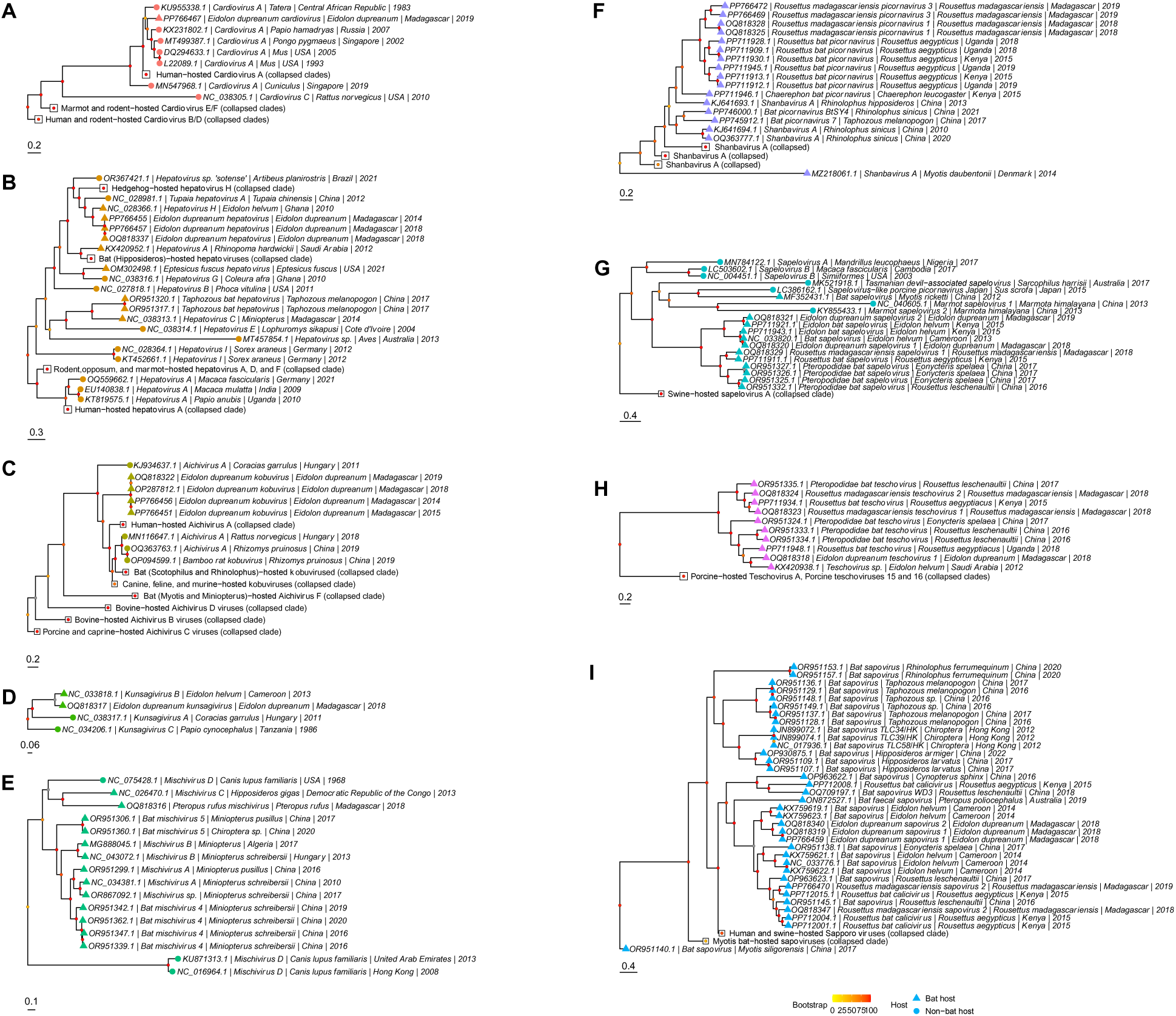
Maximum-likelihood nucleotide phylogenies (RAxML-NG-MPI^57^) of (A) *Cardiovirus* sequences across the whole genome, (B) *Hepatovirus* sequences across the whole genome, (C) *Kobuvirus* sequences across the whole genome, (D) *Kunsagivirus* sequences across the whole genome, (E) *Mischivirus* sequences across the whole genome, (F) *Shanbavirus*/unclassified bat picornavirus sequence across the whole genome, (G) *Sapelovirus* sequences across the whole genome, (H) *Teschovirus* sequences across the whole genome, and (I) *Sapovirus* sequences across the whole genome. Best fit nucleotide substitution models and overlapping base pair length per tree are summarized in **Supplemental table 1**. Bootstraps were computed using Felsenstein’s method and are visualized on tree branches. Novel sequences are highlighted in yellow. Collapsed clades are represented by white squares. Tip shape indicates host in which the sequence was derived corresponding to the legend. Trees are rooted in Sindbis virus (accession NC_001547.1). Roots have been removed for ease of visualization. Branch lengths are scaled by nucleotide substitutions per site, corresponding to each scalebar.

#### Picornaviridae

The partial *E. dupreanum cardiovirus* sequence had high identity to a divergent orangutan-hosted encephalomyocarditis virus (**Supplemental table 2**) and appeared to be phylogenetically basal to these primate-hosted cardioviruses. Due to the novelty of cardioviruses in bats, addition of more sequences would likely resolve the placement of this sequence in its own clade (**Fig. 3A**). *E. dupreanum hepatovirus* sequences formed a clade sister to a Cameroonian *E. helvum*-hosted hepatovirus (accession NC_028366), separate from insectivorous *Hipposideros* spp. bat-hosted hepatoviruses (**Fig 3B**). Consistent with previous reports^13^, human-hosted hepatoviruses form a monophyletic clade sister to a separate clade of animal-hosted hepatoviruses, and hedgehog and shrew-borne hepatoviruses form a paraphyletic clade basal to the African bat hepatoviruses (**Fig. 3B**). As mentioned previously, the *E. dupreanum kobuvirus* sequences described in this paper were determined to be genetic variants of the same virus previously described^52^. *E. dupreanum* kobuviruses were basal to other bat-hosted kobuviruses (namely *Scotophilus*/*Rhinolophus* sequences from Asia), in addition to canine and human-hosted kobuviruses, but were sister to *Aichivirus F* sequences which include *Myotis* and *Miniopterus*-hosted kobuviruses (**Fig 3C**). Few full-length *Kunsagivirus* sequences are published, but phylogenetically we can see that *E. dupreanum kunsagivirus* formed a monophyletic clade with *E. helvum kunsagivirus* B (accession NC_033818), separate from the kunsagivirus sequences found in other mammalian hosts (**Fig. 3D**). With high identity and phylogenetic clustering (**Fig. 3D**, **Supplemental table 2**), *E. dupreanum* kunsagivirus appeared to be very similar to the kunsagivirus circulating in Cameroonian *E. helvum* bats.

The most divergent virus, *P. rufus mischivirus*, formed a monophyletic clade with a *H. gigas* mischivirus C sequence from the DRC that is separate from all other bat-hosted mischiviruses that have been previously identified in *Miniopterus* spp. The long branch length separating these clades suggests considerable differences in nucleotide substitution rates as well (**Fig. 3E**). Without more sequences to resolve the phylogeny, it is unclear that geographical species distribution alone may be driving viral species differentiation. *H. gigas* is part of the family *Hipposideridae*, within the *Yinpterochiroptera* suborder (previously known as megabats), as is *P. rufus*. By contrast, *Miniopterus* spp. are part of the *Yangochiroptera* suborder (previously known as microbats)^75^. While these patterns suggest a co-speciation of viruses along host evolutionary lines, previous analyses have instead supported host-jumping mechanisms as a driver of *Mischivirus* diversity in bats^34^.

*R. madagascariensis* picornaviruses formed a sister paraphyletic clade to other *Rousettus* bat picornaviruses from East Africa, and within their own clade separated into two smaller clades representing two species (*R. madagascariensis picornavirus* 1 and 3) (**Fig. 3F**). *R. madagascariensis picornavirus* 2: accession OQ818346, was a short sequence (<2000bp in length) and was therefore excluded from phylogenetic analysis. Shanbaviruses, a *Picornaviridae* genus first described from Chinese *Miniopterus* bats, and other unclassified bat picornaviruses (*bat picornavirus* 7 and BtYS4) were basal to these African bat picornaviruses and phylogenetically distinct from a *Shanbavirus A* sequence from a Danish *Myotis daubentonii*, again suggestive of some host-virus co-speciation relationships, though further sampling will be needed to parse true evolutionary relationships (**Fig. 3F**). Bat sapeloviruses formed a monophyletic clade sister to marmot and Tasmanian-devil sapeloviruses - excepting one *Myotis*-hosted sapelovirus - with simian and porcine-hosted sapeloviruses basal in the phylogeny (**Fig. 3G**). Within the larger bat sapelovirus clade, two sister clades formed between *Eidolon*-hosted sapeloviruses and *Rousettus*/*Eonycteris*-hosted sapeloviruses (**Fig. 3G**). As seen with BLAST, *E. dupreanum sapeloviruses* 1 and 2, and *R. madagascariensis sapelovirus* 1 had respectively high identity to those viruses in Africa hosted by other bat species within the same genera (**Fig. 3G**, **Supplemental table 2**). In a similar pattern as the *Sapelovirus* phylogeny, bat teschoviruses formed a divergent and distinct clade to those hosted by swine (porcine) – which are the usual hosts of teschoviruses^76^ (**Fig. 3H**). Within the bat-hosted teschoviruses, viruses again clustered following species phylogenetics: *R. madagascariensis teschovirus* 1, *R. madagascariensis teschovirus* 2, and *E. dupreanum teschovirus* 1 grouped with similar viruses hosted by other bat species within the same respective genera (**Fig. 3H**). In general, in virus clades with prior, publicly-reported viruses for comparison, Malagasy bat *Picornaviridae* from *E. dupreanum* and *R. madagascariensis* were, respectively, phylogenetically closest to *E. helvum* and *R. aegypticus*-hosted Picornaviridae described from Cameroon, Kenya, and Uganda.

#### Caliciviridae

Bat sapoviruses, in the genus *Sapovirus*, are phylogenetically distinct from human and swine-hosted Sapporo viruses, which also fall within the *Sapovirus* genus (**Fig. 3I**). In our analyses, most bat sapoviruses formed a monophyletic clade proximal to the human and swine Sapporo viruses, *Myotis* spp. -hosted sapoviruses resolved as basal in the Sapovirus clade. This apparent paraphyly in bat Sapovirus sp. indicates a potential for host-switching events leading to the diversification of this genus (**Fig. 3I**). Novel *E. dupreanum* and *R. madagascariensis* sapoviruses again nest sister to closely related African *E. helvum* and *R. aegypticus*-hosted sapoviruses, respectively (**Fig. 3I**).

### Similarity analysis

#### Picornaviridae

Within the *Picornaviridae* family, many novel sequences had the highest identity (average >80% BLASTx^55^ identity) to African (Cameroon, Kenya, and Uganda) *E. helvum* and *R. aegypticus* viruses (**Supplemental table 2, Fig. 3A-I**), which are sister host species to *E. dupreanum* and *R. madagascariensis*. When comparing Madagascar sequences against their closest matches in GenBank, we observed a consistent sharp drop in similarity in the 2A-to-2B and 3A-to-3B broad peptide regions of the genome (**Fig. 4A-E, Supplemental fig. 3** for amino acid similarity plots**, Supplemental fig. 4** for nucleotide similarity plots). The border of the P1 and P2 regions (between VP1 and 2A, respectively) are thought to comprise a genomic region susceptible to recombination^25^, while the 3A region of *Picornaviridae*, known to be highly divergent across genera, is associated with host range determination and viral replication^77^, thus offering some explanation for the heightened genomic divergence in these regions. We also consistently observed drops in similarity in the 5’ and 3’ UTRs (**Fig. 4A-E, Supplemental fig. 3** and **4**). The 5’ UTR of *Picornaviridae* is thought to play a role in antagonizing innate host immunity; indeed, work in enteroviruses shows that the development of mutations in this region can dampen replication competence of the virus^78^. As the 5’ UTRs of most of the novel Malagasy *Picornavirales* demonstrated very low similarity (<50%) to related reference sequences (**Fig. 4A-E**), it is possible that these Malagasy bat viruses employ different replication and immune evasion strategies than previously documented for this virus family (**Fig. 4A-E**).

**Figure 4:**
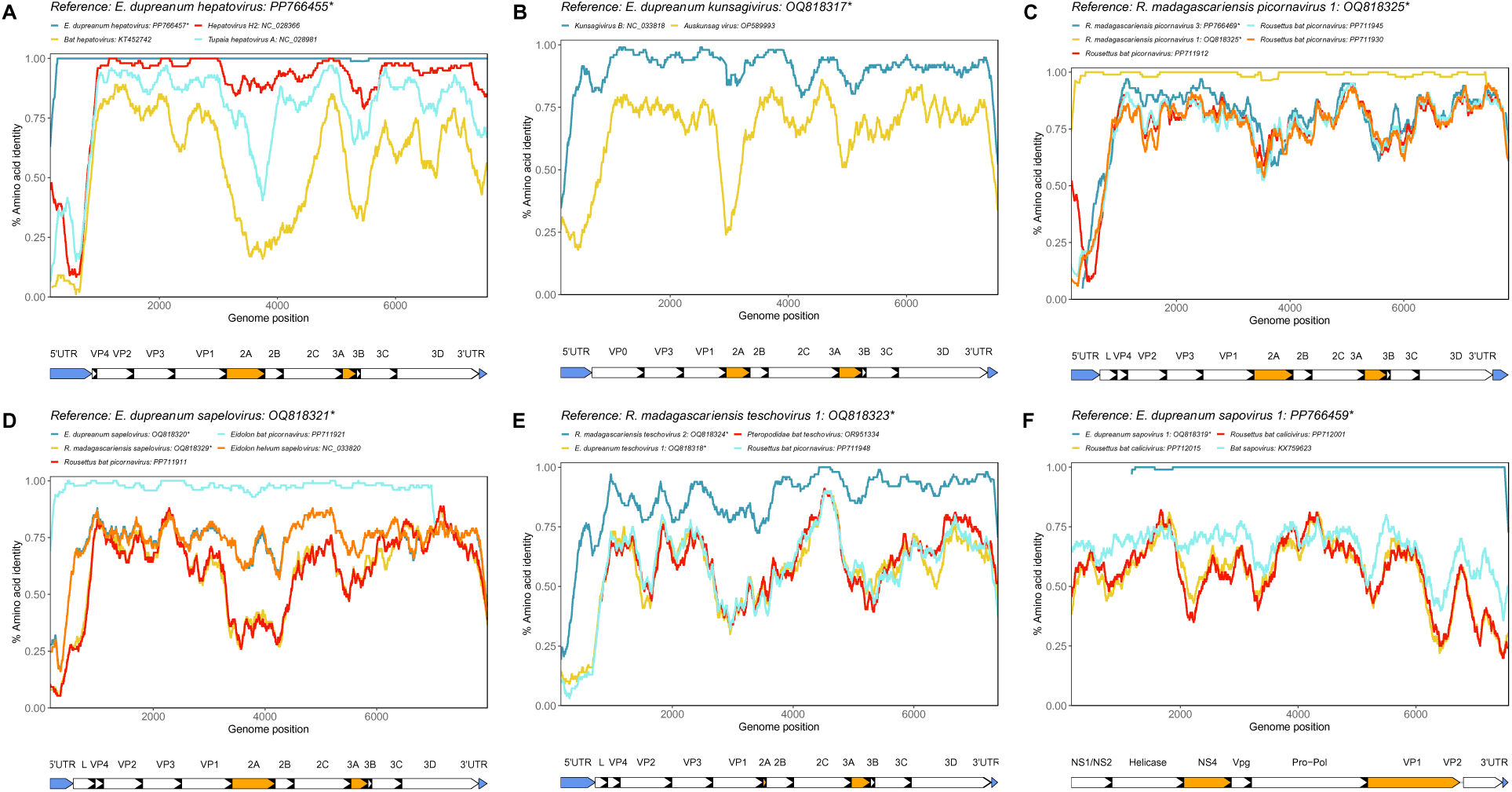
Amino acid similarity computed in PySimPlot^59^ for novel full-length sequences. Similarity analyses with query sequence (A) *E. dupreanum hepatovirus*: accession PP766455, (B) *E. dupreanum kunsagivirus*: accession OQ818217, (C) *R. madagascariensis picornavirus* 1: accession OQ818325, (D) *E. dupreanum sapelovirus*: accession OQ818321, (E) *R. madagascariensis teschovirus* 1: accession OQ818323, and (F) *E. dupreanum sapovirus* 1: accession PP766459 against similar sequences identified from BLAST and other matched novel sequences within the same genus. Novel sequences described in this study are starred with an asterisk. Line color corresponds to different virus sequences, with annotated regions of the genome below each plot. Plots were generated with a window size of 100aa and a step size of 20aa. Peptides in orange and corresponding grey shaded areas denote areas of interest for host interactions and immunogenicity, and blue peptides denote 5’ and 3’ UTRs. Amino acid similarity plots for *Kobuvirus* and *Mischivirus* are in **Supplemental fig. 3**. Matched nucleotide similarity plots are in **Supplemental fig. 4**.

For example, novel *E. dupreanum hepatovirus* sequences were nearly identical to each other, and across the genome, demonstrated highest identity to *Hepatovirus* H2, from a *E. helvum* bat (**Fig. 4A**). Nonetheless, the Malagasy viruses showed a large dip in identity to the *E. helvum* virus in the 5’ UTR (<25% average identity), in addition to other dips across the 2A/2B peptides, the 3A/3B peptides, and within the 3’ UTR (**Fig. 4A**). *E. dupreanum kunsagivirus* mirrored these patterns, showing reduced identity to previously described kunsagiviruses in the 5’ UTR, 2A, 3A, and 3’ UTR regions, in addition to a slight reduction in identity in the 2C region (**Fig. 4B**). *E. dupreanum kobuvirus* sequences described in this study were nearly identical across the genome to an *E. dupreanum kobuvirus* sequence described previously^52^ (**Supplemental fig. 3A**). *P. rufus mischivirus* had low identity to comparable sequences across the genome (<75%), with more dramatic drops observed in the 5’ UTR and the 3A peptide regions (**Supplemental fig. 3B**).

*R. madagascariensis picornavirus* 1 sequences had high (∼99%) identity across the genome to each other, and *R. madagascariensis* picornavirus 3 was most similar to African *Rousettus bat picornavirus* accession PP711945 (**Fig. 4C**). Though *R. madagascariensis picornavirus* 1 and 3 were derived from the same host species, they nonetheless demonstrated <80% average identity to one another and demonstrated predictable divergence in the 2A, 3A, and 5’ and 3’ UTR regions, suggesting that these viruses likely had different replication and immune evasion strategies (**Fig. 3C**). Similar to *R. madagascariensis* picornaviruses, the novel *E. dupreanum* and *R. madagascariensis* sapeloviruses had high identity to sister species (e.g. *E. helvum* and *R. aegypticus,* respectively)-hosted viruses, but still differed in 5’ UTR, 2A peptide, and 3A peptide regions (**Fig. 3D**). *R. madagascariensis teschovirus* 1 was not as closely related to *R. madagascariensis teschovirus* 2 (**Fig. 3E**) as were all *R. madagascariensis* bat picornaviruses (**Fig. 3C**). However, *E. dupreanum teschovirus* 1 had highest identity to *Rousettus* bat picornavirus: PP711948, derived from a *R. aegypticus* host in Uganda (**Fig. 4E**). As before, we observed drops in similarity between the Malagasy bat teschoviruses and previously described sequences in the 3A peptide region, in addition to drops in similarity in VP2 and VP1 regions, as well (**Fig. 4E**).

#### Caliciviridae

For sapoviruses, we anticipated lower similarity in the VP1, NS4, and Vpg regions of the genome. VP1 is associated with host interactions and immunogenicity^79^, while *Caliciviridae* NS4 has been suggested to be a homolog for *Picornaviridae* 3A, since both families exhibit similar genome organization^80^. If true, then divergence in NS4 might be expected due to divergence in host range, though the function of NS4 is not well characterized. Vpg (or NS5 in some sources), is a nonstructural protein that primes genome replication^81^.

Both novel *E. dupreanum sapovirus* 1 sequences were nearly identical to each other, but *E. dupreanum sapovirus* 1: accession OQ818319 is a partial genome and was missing the NS1/NS2 region of the genome, so it is possible some variation may exist in this region in the two sequences (**Fig. 4F**). The most similar virus, a Cameroonian *E. helvum sapovirus*, hovered around 70% identity to the novel query sequences across most of the genome, with three drops in similarity at the border of Vpg and Pro-Pol, on the border of Pro-Pol and VP1, and in VP1 (**Fig. 4F**). The drops in identity were not as dramatic as those witnessed in certain regions for other novel Malagasy bat picornaviruses; in general, *E. dupreanum sapovirus* 1 displayed an average around 50% identity to previously characterized viruses in this clade (**Fig. 4F**). As observed in Malagasy bat *Picornaviridae* (**Fig. 4A-E**), dips in identity across the genome corresponding to areas involving host interactions and immune responses could indicate that while *E. dupreanum* and *R. madagascariensis*-hosted sapoviruses were respectively most similar to African *E. helvum* and *R. aegypticu*s-hosted sapoviruses (range from ∼67% to ∼90% identity) (**Supplemental table 3**), these novel viruses likely use different replication and immune evasion strategies. All R. madagascariensis sapoviruses were partial genomes, so were not included in genome wide similarity analysis or subsequent recombination analysis.

### RDP4 recombination analysis

#### Picornaviridae

Recombination analysis performed on novel Malagasy *Picornaviridae* indicated that there is evidence for genetic exchange with viruses hosted by African *E. helvum* and *R. aegypticus* (**Fig. 5**, **Supplemental table 6**). Analyses using full genome novel *Picornaviridae* and high identity reference sequences identified from BLAST^55^ (**Supplemental table 2**), phylogenetic analysis (**Fig. 2 and Fig. 3A-I**), and genome-wide similarity scans (**Fig. 4**), indicated that the following genera were most likely to be under recombination pressure: *Hepatovirus*, *Sapelovirus*, and *Teschovirus* (**Supplemental table 5**). No significant recombination pressure was observed in *Cardiovirus*, *Kunsagivirus*, *Kobuvirus*, and *Mischivirus* (**Supplemental table 6**). Some recombination pressure was observed in *R. madagascariensis* picornaviruses and *Sapovirus* but was not further analyzed due to either fewer than four significant tests for recombination and/or no novel sequence identified as a recombinant sequence or major parent sequence in the alignment (**Supplemental table 6**).

**Figure 5:**
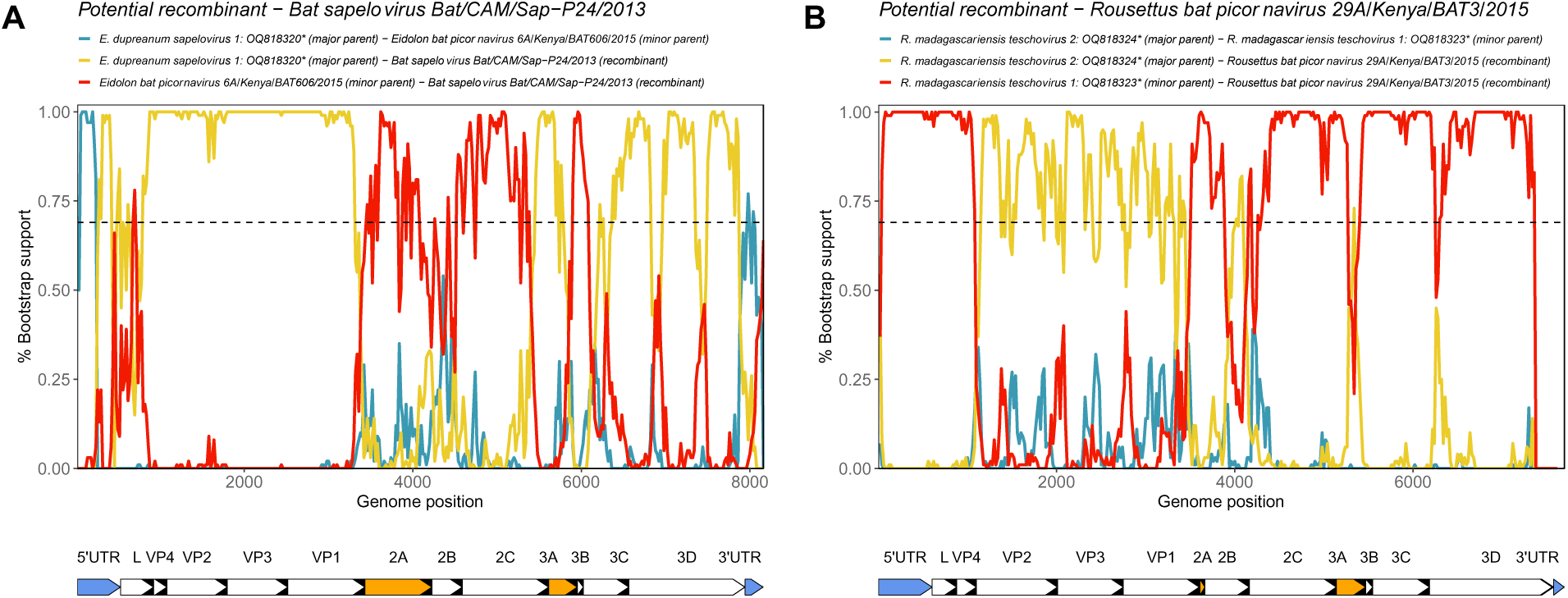
Bootscan plots computed in RDP4^60^ for potential recombinant sequences (A) *Bat sapelovirus* Bat/CAM/Sap-p24/2013: accession NC_033820 and (B) *Rousettus bat picornavirus* 29A/Kenya/BAT3/2015: accession PP711934. Line color corresponds to pairwise alignments between the potential recombinant sequence, major parental sequence, and the minor parental sequence. Asterisks denote novel sequences described in this study. Horizontal dashed line refers to a 70% cutoff bootstrap percentage, and grey bars indicate regions identified as significant areas of recombination (P<0.05) across at least 5 analyses within RDP4^60^ (RDP, GENECONV, Bootscan, Maxchi, Chimaera, and 3Seq). Nucleotide bootscan plots were generated using a window size of 200bp and a step size of 20bp. Genome maps are below each plot, peptides in orange denote areas of interest for host interactions and immunogenicity, and blue peptides denote 5’ and 3’ UTRs. RDP4^60^ statistics are reported in **Supplemental table 6**.

Through RDP4 analysis, *E. dupreanum hepatoviruses* (both accession PP766455 and PP766457) were identified as possible minor parental sequences to recombinant *E. helvum hepatovirus* M32EidHel2010 (accession NC_028366), with breakpoints across the genome; however, bootstrap support was too weak to indicate any other parental sources of genomic material (**Supplemental fig. 5A**). Within *Sapelovirus*, the strongest bootstrap support indicated that Cameroonian *bat sapelovirus* Bat/CAM/Sap-P24/2013 (accession NC_033820) was the likely recombinant sequence with genomic material from *E. dupreanum sapelovirus* 1 (major parental sequence) (**Fig. 5A**). The highest support across most of the genome was for *E. dupreanum sapelovirus* 1 as a major parental sequence excepting the P2 region (2A, 2B, and 2C) and the 3A peptide where higher bootstrap support indicated that Kenyan *Eidolon bat picornavirus* 6A/Kenya/BAT606/2015 was the minor parental sequence (accession PP711943) (**Fig. 5A**). There was additional evidence that *E. dupreanum sapelovirus* 2 was a recombinant sequence with Kenyan *Eidolon bat picornavirus* 6A/Kenya/BAT606/2015 as a minor parental sequence across most of the genome, with highest bootstrap support for this genomic history in the P2 (2A, 2B, 2C) region and lower bootstrap support in P1 region (VP4, VP2, VP3, and VP1) (**Supplemental fig. 5B**).

Evidence for recombination was highest overall within the *Teschovirus* genus (**Fig. 5B** and **Supplemental fig. 5C** and **5D**), where the most likely recombinant was identified as Kenyan *Rousettus bat picornavirus* 29A/Kenya/BAT3/2015 (accession PP711934), with genomic material contributed from *R. madagascariensis teschovirus* 2 (major parental sequence) and *R. madagascariensis teschovirus* 1 (minor parental sequence), with higher support for the major parental sequence in the P1 region (VP4, VP2, VP3, and VP1) and higher support for the minor parental sequence among the rest of the genome (**Fig. 5B**). Additional analysis with a consensus sequence of *R. aegypticus teschovirus* sequences (accessions PP711948 and PP711934) further supported the involvement of *R. madagascariensis teschovirus* 1 and 2 in recombination, but with lower bootstrap support across the whole genome (**Supplemental fig. 5C** and **5D**). Support for *R. madagascariensis teschovirus* 1 as a recombinant sequence with *R. aegypticus teschovirus* clade as a minor parental sequence was highest in the 5’UTR/3D region (**Supplemental fig. 5C**). The opposite pattern was observed in the 3’UTR in analysis of *R. aegypticus teschovirus* clade as a recombinant sequence with *R. madagascariensis teschovirus* 1 as a major parental sequence (**Supplemental fig. 5D**). VP1-VP3 peptides in *Picornaviridae* are thought to play a role in generation of neutralizing antibodies^82^, and as previously mentioned, the P1-P2 junction, 3A/3B peptides, and 5’ UTR also play a role in determination of host range and immune response^25,77,78^.

## DISCUSSION

Using mNGS data, we analyzed the diversity of *Picornavirales*, namely *Picornaviridae* and *Caliciviridae*, finding evidence of high viral similarity between sequences from Malagasy fruit bats (*E. dupreanum* and *R. madagascariensis*) with those from their sister species (*E. helvum* and *R. aegypticus*, respectively) in the mainland African countries of Cameroon, Kenya, and Uganda, although it is possible these countries simply represent the most intensely sampled localities to date. Further, we find evidence that recombination between these similar sequences may have led to the diversification of these viral genera in Malagasy fruit bats through phylogenetic clustering and recombination analyses. We have previously demonstrated potential cross-continental viral genetic exchange between African and Malagasy fruit bat-hosted nobecoviruses, so it is possible these *Picornaviridae* and *Caliciviridae* are diversifying in similar manners^46^.

We identified 13 full-length and 37 partial-length (ranging from 1014 to 8379 bp in length) genomic sequences (14 *Caliciviridae* and 36 *Picornaviridae* sequences) from Malagasy fruit bats: *E. dupreanum*, *P. rufus*, and *R. madagascariensis*. Multiple *Picornaviridae* and *Caliciviridae* species were identified within the same sampling dates at *E. dupreanum* and *R. madagascariensis* roost sites, suggesting that many of these diverse viruses shed simultaneously within a representative bat population, a pattern that has been reported before for fruit bats^83^. Bats captured at a site within the same sampling session could reflect that roost’s viral population dynamics at a single timepoint, as they often shared similar virome profiles. Indeed, viral diversity was significantly different between roost sites dominated by *E. dupreanum* versus *R. madagascariensis*, and sapeloviruses, teschoviruses, and sapoviruses all resolved into disparate host species-specific clades. *R. madagascariensis* picornaviruses, which displayed the lowest identity to previously-known sequences (∼80% to known Ugandan/Kenyan *R. aegypticus*-hosted picornaviruses^63^), formed their own clade distinct from their *Rousettus*-hosted viruses described elsewhere, which further separated into two separate clades corresponding to two different viral species.

Many of the novel sequences we identified in this analysis displayed reduced identity to previously-described sequences in genomic areas that determine host range and immune response (5’ UTR, 2A/2B peptides, and 3A/3B peptides in *Picornaviridae*^25,77,78^, NS4 and VP1 peptides in *Caliciviridae*^80,81^). These regions of reduced similarity indicate that, despite relatedness to sister species-hosted viruses in Africa, Malagasy bat-hosted viruses could employ different replication and immune evasion mechanisms. Our recombination analysis did not identify the 3A region, which is highly diverse in *Picornaviridae* and likely contributes to host range^77^, to be under recombination pressure. To our knowledge, recombination analysis has not been previously performed on any African bat *Picornavirales* sequences^35,36,63,84^. While no evidence for local recombination between the novel Malagasy viruses was found, analyses presented here suggest that recombination likely contributed to the diversification of this clade in bat hosts, and in some cases may have predated dispersal of these viruses between Madagascar and mainland Africa. Of note, we observed evidence of more virus genetic exchange between *E. dupreanum* and *R. madagascariensis* vs. *P. rufus*. The former two bat species are known to co-roost in caves in our system^46^, while the latter species is tree-dwelling, highlighting the importance of host proximity in viral dispersal, exchange, and diversification. Cave co-roosting has been previously shown to support recombination and diversification in bat coronavirus systems^85^. As *E. dupreanum* and *R. madagascariensis* are known to co-roost with insectivorous bats on occasion^86^, additional screening for these viral families in more Malagasy bat species may further support evidence for high fidelity to a single host species.

In Madagascar and some areas of mainland Africa, humans hunt bats for food^87–89^. While no bat-borne zoonosis has been linked to this practice in Madagascar, undiagnosed fevers are common, and it is possible that bat virus zoonoses may be occurring undetected^90,91^. Enteric viruses described from Cameroonian hunters demonstrated a high diversity of *Picornavirales*, including some sequences which share evolutionary ancestry with bat- or other animal-hosted viruses^84^. Nonetheless, human-hosted *Picornavirales* in Cameroon segregate phylogenetically from animal viruses, suggesting that, while zoonosis may be possible in this clade, these cross-species emergence events are relatively rare^84^. Indeed, discrete host-species relationships appear to drive most of the observed diversification within the *Picornavirales* clade. While we lack *Picornavirales* sequences from humans in Madagascar to test these hypotheses, the bat sequences described here group with other animal (particularly bat)-derived viruses from related host species^63^. As recombination events can precede more dramatic host switches, including zoonoses, understanding of these processes within diverse viral clades is critical to assessing potential zoonotic risk.

## ETHICS STATEMENT

The animal study was reviewed and approved by UC Berkeley Animal Care and Use Committee and Madagascar Ministry of Forest and the Environment under guidelines posted by the American Veterinary Medical Association.

## ACKNOWLEDGMENTS

The authors acknowledge Anecia Gentles, Kimberly Rivera, Fifi Ravelomanantsoa, and Sarah Guth for help in the field and lab. We acknowledge the Virology Unit at the Institut Pasteur de Madagascar. We thank Amy Kistler, Cristina M. Tato, Maira Phelps, Vida Ahyong, Angela Detweiler, Michelle Tan, Norma Neff, and Joseph L. DeRisi of the Chan Zuckerberg Biohub (CZB) for logistical support. We additionally thank Angela Detweiler, Michelle Tan, and Norma Neff of the CZB genomics platform for mNGS support and thank the Brook lab at the University of Chicago for helpful contributions to the manuscript. This work was completed in part with resources provided by the University of Chicago’s Research Computing Center.

## DATA AVAILABILITY

All full and partial length genome sequences were submitted to NCBI and assigned accession numbers OQ818316-OQ818318, OQ818320-OQ818324, OQ818328, OQ818329, PP766456, PP766459, PP766469 (full-length genomes), and OQ818319, OQ818325, OQ818337, OQ818340, OQ818342-OQ818348, PP766449-PP766455, PP766457, PP766458, PP766460-PP766468, PP766470-PP766477 (partial-length genomes). Detailed descriptions of analyses done in this paper, including scripts used to generate figures, are available on our GitHub (https://github.com/brooklabteam/mada-bat-picornavirus).

## AUTHOR CONTRIBUTIONS

CEB conceived of the project and acquired the funding, in collaboration with J-MH, VL, and PD. Field samples were collected, and RNA extracted by CEB, HCR, SA, AA, TR, GK, and VR. AK led the mNGS at CZB, with support from VA, HCR, TR, CEB, and JLD. ARH led the mNGS at NIH, with support from FL, LL, RD, SG, CEB, and DCD. GK analyzed the resulting data and wrote the original draft of the manuscript with CEB, which all authors edited and approved. SH and ECR provided noteworthy edits to the manuscript and contributed to finalization of the manuscript.

## CITATIONS

1. Le Gall, O., et al. Picornavirales, a proposed order of positive-sense single-stranded RNA viruses with a pseudo-T = 3 virion architecture. Arch. Virol. 153, 715 (2008).

2. Cao, G., Jing, W., Liu, J. & Liu, M. The global trends and regional differences in incidence and mortality of hepatitis A from 1990 to 2019 and implications for its prevention. Hepatol. Int. 15, 1068–1082 (2021).

3. Mehndiratta, M. M., Mehndiratta, P. & Pande, R. Poliomyelitis: Historical Facts, Epidemiology, and Current Challenges in Eradication. The Neurohospitalist 4, 223–229 (2014).

4. Human Viruses: Diseases, Treatments and Vaccines: The New Insights. (Springer International Publishing, Cham, 2021). doi:10.1007/978-3-030-71165-8.

5. De Graaf, M., Van Beek, J. & Koopmans, M. P. G. Human norovirus transmission and evolution in a changing world. Nat. Rev. Microbiol. 14, 421–433 (2016).

6. Becker-Dreps, S., Bucardo, F. & Vinjé, J. Sapovirus: an important cause of acute gastroenteritis in children. Lancet Child Adolesc. Health 3, 758–759 (2019).

7. Olival, K. J., et al. Host and viral traits predict zoonotic spillover from mammals. Nature 546, 646–650 (2017).

8. Mollentze, N. & Streicker, D. G. Viral zoonotic risk is homogenous among taxonomic orders of mammalian and avian reservoir hosts. Proc. Natl. Acad. Sci. 117, 9423–9430 (2020).

9. Bailey, E. S., Fieldhouse, J. K., Choi, J. Y. & Gray, G. C. A Mini Review of the Zoonotic Threat Potential of Influenza Viruses, Coronaviruses, Adenoviruses, and Enteroviruses. Front. Public Health 6, 104 (2018).

10. Desselberger, U. Caliciviridae Other Than Noroviruses. Viruses 11, 286 (2019).

11. Ji, C., et al. Systematic Surveillance of an Emerging Picornavirus among Cattle and Sheep in China. Microbiol. Spectr. 11, e05040–22 (2023).

12. Cui, X., et al. Virus diversity, wildlife-domestic animal circulation and potential zoonotic viruses of small mammals, pangolins and zoo animals. Nat. Commun. 14, 2488 (2023).

13. Drexler, J. F., et al. Evolutionary origins of hepatitis A virus in small mammals. Proc. Natl. Acad. Sci. 112, 15190–15195 (2015).

14. Kocher, J. F., et al. Bat Caliciviruses and Human Noroviruses Are Antigenically Similar and Have Overlapping Histo-Blood Group Antigen Binding Profiles. mBio 9, e00869–18 (2018).

15. Jamal, S. M. & Belsham, G. J. Foot-and-mouth disease: past, present and future. Vet. Res. 44, 116 (2013).

16. Tan, S. Z. K., Tan, M. Z. Y. & Prabakaran, M. Saffold virus, an emerging human cardiovirus: Pathogenesis of Saffold virus. Rev. Med. Virol. 27, e1908 (2017).

17. Mattison, K., et al. Human Noroviruses in Swine and Cattle. Emerg. Infect. Dis. 13, 1184–1188 (2007).

18. Harvala, H., et al. Co-circulation of enteroviruses between apes and humans. J. Gen. Virol. 95, 403–407 (2014).

19. Harvala, H., et al. Detection and Genetic Characterization of Enteroviruses Circulating among Wild Populations of Chimpanzees in Cameroon: Relationship with Human and Simian Enteroviruses. J. Virol. 85, 4480–4486 (2011).

20. Coyne, K. P., et al. Recombination of Feline calicivirus within an endemically infected cat colony. J. Gen. Virol. 87, 921–926 (2006).

21. Hansman, G. S., et al. Intergenogroup Recombination in Sapoviruses. Emerg. Infect. Dis. 11, 1914–1920 (2005).

22. Heath, L., Van Der Walt, E., Varsani, A. & Martin, D. P. Recombination Patterns in Aphthoviruses Mirror Those Found in Other Picornaviruses. J. Virol. 80, 11827–11832 (2006).

23. Kempf, B. J., Watkins, C. L., Peersen, O. B. & Barton, D. J. Picornavirus RNA Recombination Counteracts Error Catastrophe. J. Virol. 93, e00652–19 (2019).

24. Tolf, C., et al. Molecular characterization of a novel Ljungan virus (Parechovirus; Picornaviridae) reveals a fourth genotype and indicates ancestral recombination. J. Gen. Virol. 90, 843–853 (2009).

25. Lukashev, A. N. Recombination among picornaviruses: Recombination among picornaviruses. Rev. Med. Virol. 20, 327–337 (2010).

26. Zhou, P., et al. Fatal swine acute diarrhoea syndrome caused by an HKU2-related coronavirus of bat origin. Nature 556, 255–258 (2018).

27. Jiang, S., Xia, S., Ying, T. & Lu, L. A novel coronavirus (2019-nCoV) causing pneumonia-associated respiratory syndrome. Cell. Mol. Immunol. 17, 554–554 (2020).

28. Memish, Z. A., et al. Middle East Respiratory Syndrome Coronavirus in Bats, Saudi Arabia. Emerg. Infect. Dis. 19, (2013).

29. Leendertz, S. A. J., Gogarten, J. F., Düx, A., Calvignac-Spencer, S. & Leendertz, F. H. Assessing the Evidence Supporting Fruit Bats as the Primary Reservoirs for Ebola Viruses. EcoHealth 13, 18–25 (2016).

30. Guth, S., et al. Bats host the most virulent—but not the most dangerous—zoonotic viruses. Proc. Natl. Acad. Sci. 119, e2113628119 (2022).

31. Schountz, T., Baker, M. L., Butler, J. & Munster, V. Immunological Control of Viral Infections in Bats and the Emergence of Viruses Highly Pathogenic to Humans. Front. Immunol. 8, 1098 (2017).

32. Diakoudi, G., et al. Genome sequence of an aichivirus detected in a common pipistrelle bat (Pipistrellus pipistrellus). Arch. Virol. 165, 1019–1022 (2020).

33. Kemenesi, G., et al. Genetic characterization of a novel picornavirus detected in Miniopterus schreibersii bats. J. Gen. Virol. 96, 815–821 (2015).

34. Zeghbib, S., et al. Genetic characterization of a novel picornavirus in Algerian bats: co-evolution analysis of bat-related picornaviruses. Sci. Rep. 9, 15706 (2019).

35. Yinda, C. K., et al. Novel highly divergent sapoviruses detected by metagenomics analysis in straw-colored fruit bats in Cameroon: Divergent bat sapoviruses. Emerg. Microbes Infect. 6, 1–7 (2017).

36. Yinda, C. K., et al. Highly diverse population of Picornaviridae and other members of the Picornavirales, in Cameroonian fruit bats. BMC Genomics 18, 249 (2017).

37. Waruhiu, C., et al. Molecular detection of viruses in Kenyan bats and discovery ofnovel astroviruses, caliciviruses and rotaviruses. Virol. Sin. 32, 101–114 (2017).

38. Wang, J., et al. Discovery of novel virus sequences in an isolated and threatened bat species, the New Zealand lesser short-tailed bat (Mystacina tuberculata). J. Gen. Virol. 96, 2442–2452 (2015).

39. Tse, H., et al. Discovery and Genomic Characterization of a Novel Bat Sapovirus with Unusual Genomic Features and Phylogenetic Position. PLoS ONE 7, e34987 (2012).

40. Firth, C., et al. Detection of Zoonotic Pathogens and Characterization of Novel Viruses Carried by Commensal Rattus norvegicus in New York City. mBio 5, e01933–14 (2014).

41. Scheuer, K. A., et al. Prevalence of Porcine Noroviruses, Molecular Characterization of Emerging Porcine Sapoviruses from Finisher Swine in the United States, and Unified Classification Scheme for Sapoviruses. J. Clin. Microbiol. 51, 2344–2353 (2013).

42. Martella, V., et al. Identification of a Porcine Calicivirus Related Genetically to Human Sapoviruses. J. Clin. Microbiol. 46, 1907–1913 (2008).

43. Mombo, I. M., et al. Characterization of a Genogroup I Sapovirus Isolated from Chimpanzees in the Republic of Congo. Genome Announc. 2, e00680–14 (2014).

44. Lewis-Rogers, N. & Crandall, K. A. Evolution of Picornaviridae: An examination of phylogenetic relationships and cophylogeny. Mol. Phylogenet. Evol. 54, 995–1005 (2010).

45. Antonelli, A., et al. Madagascar’s extraordinary biodiversity: Evolution, distribution, and use. Science 378, eabf0869 (2022).

46. Kettenburg, G., et al. Full Genome Nobecovirus Sequences From Malagasy Fruit Bats Define a Unique Evolutionary History for This Coronavirus Clade. Front. Public Health 10, 786060 (2022).

47. Madera, S., et al. Discovery and Genomic Characterization of a Novel Henipavirus, Angavokely Virus, from Fruit Bats in Madagascar. J. Virol. 96, e00921–22 (2022).

48. Razanajatovo, N. H., et al. Detection of new genetic variants of Betacoronaviruses in Endemic Frugivorous Bats of Madagascar. Virol. J. 12, 42 (2015).

49. Mélade, J., et al. An eco-epidemiological study of Morbilli-related paramyxovirus infection in Madagascar bats reveals host-switching as the dominant macro-evolutionary mechanism. Sci. Rep. 6, 23752 (2016).

50. Joffrin, L., et al. Bat coronavirus phylogeography in the Western Indian Ocean. Sci. Rep. 10, 6873 (2020).

51. Horigan, S., et al. Detection, characterization, and phylogenetic analysis of novel astroviruses from endemic Malagasy fruit bats. Virol. J. 21, 195 (2024).

52. Gonzalez, F. L., et al. Genomic characterization of novel bat kobuviruses in Madagascar: implications for viral evolution and zoonotic risk. Preprint at 10.1101/2024.12.24.630179 (2024).

53. Brook, C. E., et al. Disentangling serology to elucidate henipa- and filovirus transmission in Madagascar fruit bats. J. Anim. Ecol. 88, 1001–1016 (2019).

54. Kalantar, K. L., et al. IDseq—An open source cloud-based pipeline and analysis service for metagenomic pathogen detection and monitoring. GigaScience 9, giaa111 (2020).

55. BLAST® Command Line Applications User Manual.

56. Katoh, K. MAFFT: a novel method for rapid multiple sequence alignment based on fast Fourier transform. Nucleic Acids Res. 30, 3059–3066 (2002).

57. Stamatakis, A. RAxML version 8: a tool for phylogenetic analysis and post-analysis of large phylogenies. Bioinformatics 30, 1312–1313 (2014).

58. Darriba. modeltest. GitHub https://github.com/ddarriba/modeltest.

59. Davies. PySimPlot. GitHub https://github.com/jonathanrd/PySimPlot.

60. Martin, D. P., Murrell, B., Golden, M., Khoosal, A. & Muhire, B. RDP4: Detection and analysis of recombination patterns in virus genomes. Virus Evol. 1, vev003 (2015).

61. Shi, J. J., et al. A Deep Divergence Time between Sister Species of *Eidolon* (Pteropodidae) with Evidence for Widespread Panmixia. Acta Chiropterologica 16, 279–292 (2014).

62. Yeo, D. S.-Y., et al. A highly divergent Encephalomyocarditis virus isolated from nonhuman primates in Singapore. Virol. J. 10, 248 (2013).

63. Wang, D., et al. Substantial viral diversity in bats and rodents from East Africa: insights into evolution, recombination, and cocirculation. Microbiome 12, 72 (2024).

64. Doysabas, K. C. C., et al. Encephalomyocarditis virus is potentially derived from eastern bent-wing bats living in East Asian countries. Virus Res. 259, 62–67 (2019).

65. Mishra, N., et al. A viral metagenomic survey identifies known and novel mammalian viruses in bats from Saudi Arabia. PLOS ONE 14, e0214227 (2019).

66. Yinda, C. K., et al. Novel highly divergent reassortant bat rotaviruses in Cameroon, without evidence of zoonosis. Sci. Rep. 6, 34209 (2016).

67. Goodman, S. M., Chan, L. M., Nowak, M. D. & Yoder, A. D. Phylogeny and biogeography of western Indian Ocean *Rousettus* (Chiroptera: Pteropodidae). J. Mammal. 91, 593–606 (2010).

68. Lau, S. K. P., et al. Complete Genome Analysis of Three Novel Picornaviruses from Diverse Bat Species. J. Virol. 85, 8819–8828 (2011).

69. Wang, J., et al. Individual bat virome analysis reveals co-infection and spillover among bats and virus zoonotic potential. Nat. Commun. 14, 4079 (2023).

70. Mohamed, F. F., et al. Detection and genetic characterization of bovine kobuvirus from calves in Egypt. Arch. Virol. 163, 1439–1447 (2018).

71. Olarte-Castillo, X. A., et al. Molecular characterization of canine kobuvirus in wild carnivores and the domestic dog in Africa. Virology 477, 89–97 (2015).

72. Wu, Z., et al. Deciphering the bat virome catalog to better understand the ecological diversity of bat viruses and the bat origin of emerging infectious diseases. ISME J. 10, 609–620 (2016).

73. Altan, E., et al. Enteric virome of Ethiopian children participating in a clean water intervention trial. PLOS ONE 13, e0202054 (2018).

74. Joffret, M. L., et al. Genomic characterization of Sebokele virus 1 (SEBV1) reveals a new candidate species among the genus Parechovirus. J. Gen. Virol. 94, 1547–1553 (2013).

75. Lei, M. & Dong, D. Phylogenomic analyses of bat subordinal relationships based on transcriptome data. Sci. Rep. 6, 27726 (2016).

76. Malik, Y. S. et al. Teschovirus. in Emerging and Transboundary Animal Viruses (eds. Malik, Y. S., Singh, R. K. & Yadav, M. P.) 123–136 (Springer Singapore, Singapore, 2020). doi:10.1007/978-981-15-0402-0_6.

77. Jackson, T. & Belsham, G. J. Picornaviruses: A View from 3A. Viruses 13, 456 (2021).

78. Kloc, A., Rai, D. K. & Rieder, E. The Roles of Picornavirus Untranslated Regions in Infection and Innate Immunity. Front. Microbiol. 9, 485 (2018).

79. Miyazaki, N., et al. Atomic Structure of the Human Sapovirus Capsid Reveals a Unique Capsid Protein Conformation in Caliciviruses. J. Virol. 96, e00298–22 (2022).

80. Smertina, E., Hall, R. N., Urakova, N., Strive, T. & Frese, M. Calicivirus Non-structural Proteins: Potential Functions in Replication and Host Cell Manipulation. Front. Microbiol. 12, 712710 (2021).

81. Ji, X., et al. Genomic Characterization and Molecular Evolution of Sapovirus in Children under 5 Years of Age. Viruses 16, 146 (2024).

82. McKnight, K. L. & Lemon, S. M. Hepatitis A Virus Genome Organization and Replication Strategy. Cold Spring Harb. Perspect. Med. 8, a033480 (2018).

83. Peel, A. J., et al. Synchronous shedding of multiple bat paramyxoviruses coincides with peak periods of Hendra virus spillover. Emerg. Microbes Infect. 8, 1314–1323 (2019).

84. Yinda, C. K., et al. Gut Virome Analysis of Cameroonians Reveals High Diversity of Enteric Viruses, Including Potential Interspecies Transmitted Viruses. mSphere 4, e00585–18 (2019).

85. Hu, B. et al. Discovery of a rich gene pool of bat SARS-related coronaviruses provides new insights into the origin of SARS coronavirus. PLOS Pathog. 13, e1006698 (2017).

86. Andrianiaina, A., et al. Reproduction, seasonal morphology, and juvenile growth in three Malagasy fruit bats. J. Mammal. 103, 1397–1408 (2022).

87. Cardiff, S. G. & Jenkins, R. K. B. The Bats of Madagascar: A Conservation Challenge. (2016).

88. Jenkins, R. & Racey, P. Bats as bushmeat in Madagascar. Madag. Conserv. Dev. 3, (2009).

89. Cardiff, S. G., Ratrimomanarivo, F. H., Rembert, G. & Goodman, S. M. Hunting, disturbance and roost persistence of bats in caves at Ankarana, northern Madagascar. Afr. J. Ecol. 47, 640–649 (2009).

90. Mathiot, C. C., Fontenille, D., Georges, A. J. & Coulanges, P. Antibodies to haemorrhagic fever viruses in Madagascar populations. Trans. R. Soc. Trop. Med. Hyg. 83, 407–409 (1989).

91. Guillebaud, J., et al. Study on causes of fever in primary healthcare center uncovers pathogens of public health concern in Madagascar. PLoS Negl. Trop. Dis. 12, e0006642 (2018).

92. Fu, L., Niu, B., Zhu, Z., Wu, S. & Li, W. CD-HIT: accelerated for clustering the next-generation sequencing data. Bioinformatics 28, 3150–3152 (2012).

