## Supplemental figures for "*Picornaviridae* and *Caliciviridae* diversity in Madagascar fruit bats is driven by cross-continental genetic exchange"

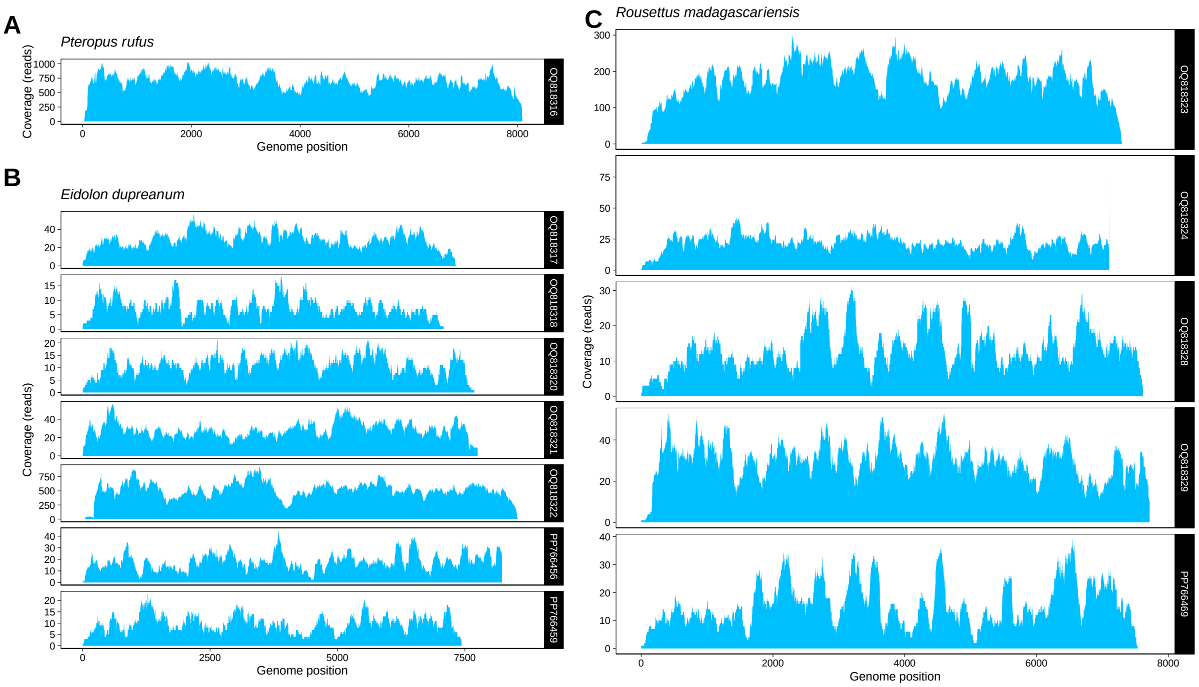


**Supplemental figure 1**: Read depth and coverage after deduplication by CD-HIT[92] for full-length contigs assembled in CZID recovered from (A) *P. rufus*, (B) *E. dupreanum*, and (C) *R. madagascariensis*. Contig depth is shown in the plots as raw coverage (number of reads mapped to genome at each nucleotide position) recovered for each full genome sequence.

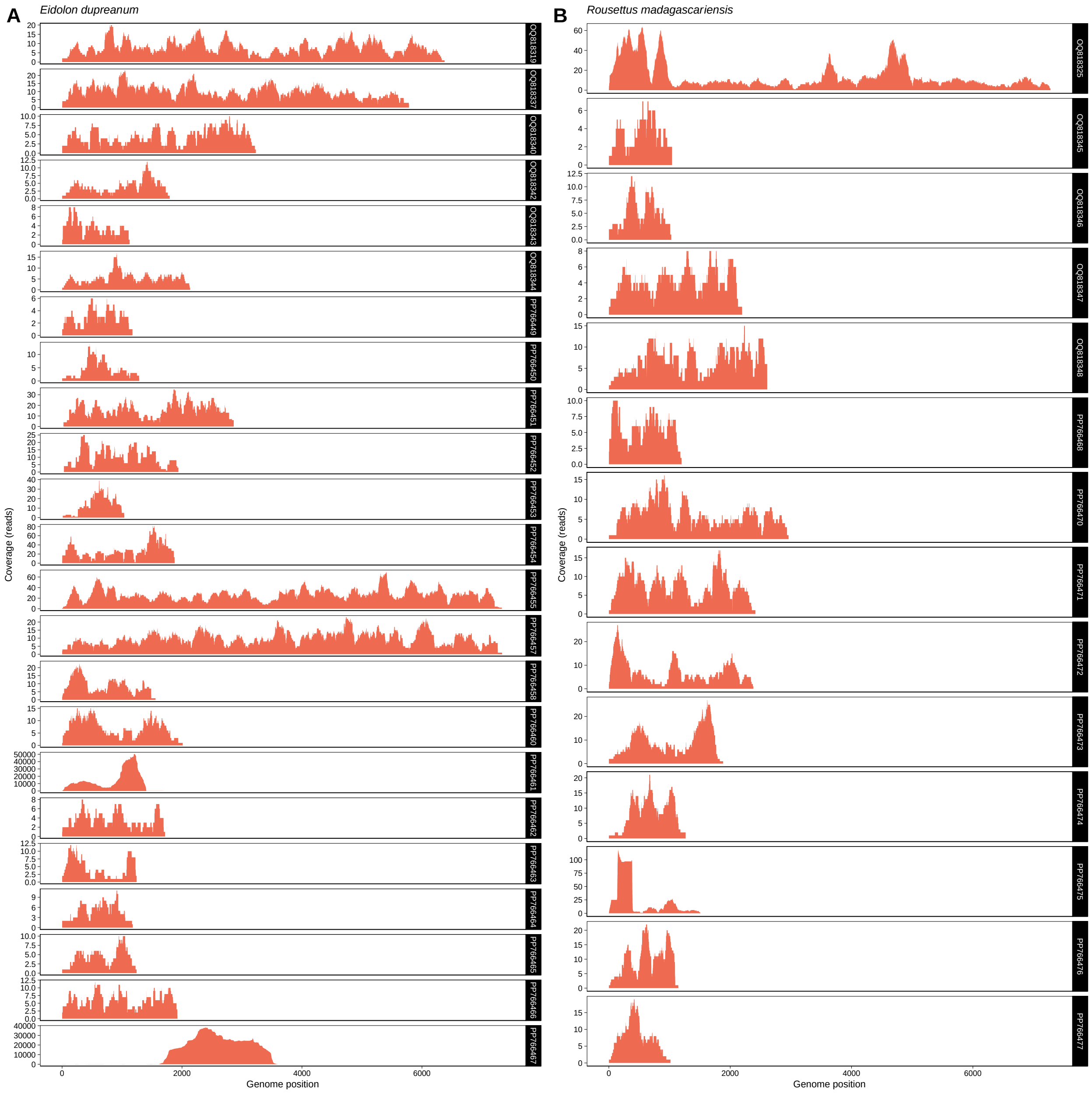


**Supplemental figure 2**: Read depth and coverage after deduplication by CD-HIT[92] for partial-length contigs assembled in CZID recovered from (A) *E. dupreanum* and (B) *R. madagascariensis*. Contig depth is shown in the plots as raw coverage (number of reads mapped to genome at each nucleotide position) recovered for each partial genome sequence.

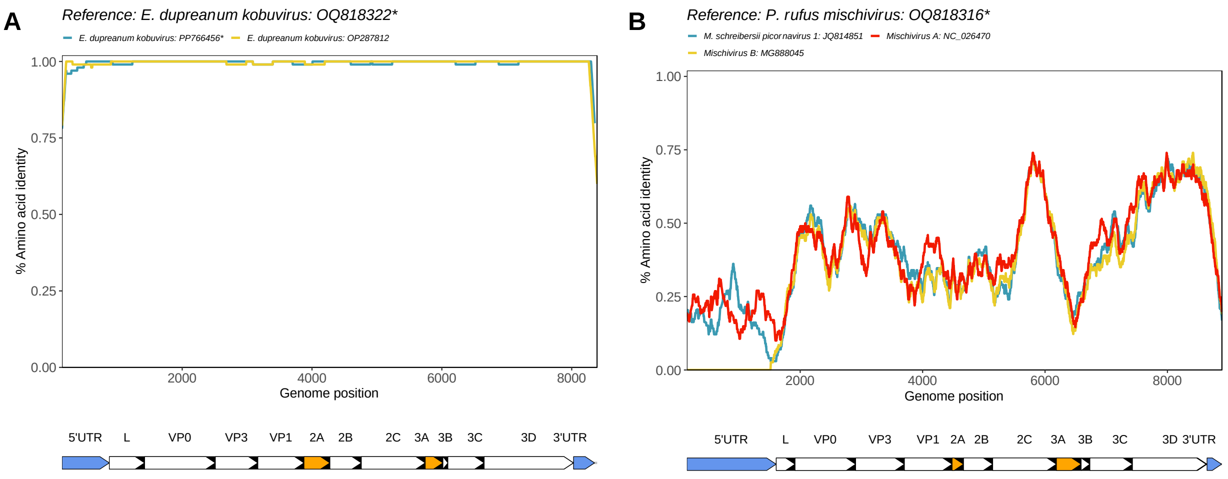


**Supplemental figure 3**: Amino acid similarity computed in PySimPlot[59] for novel full-length sequences. Similarity analyses with query sequence (A) *E. dupreanum kobuvirus*: accession OQ818322 and (B) *P. rufus mischivirus*: accession OQ818316 against similar sequences identified from BLAST[55] and other matched novel sequences within the same genus. Asterisks denote novel sequences described in this study. Line color corresponds to different virus sequences, with annotated regions of the genome below each plot. Peptides in orange and corresponding grey shaded areas denote areas of interest for host interactions and immunogenicity, and blue peptides denote 5' and 3' UTRs.

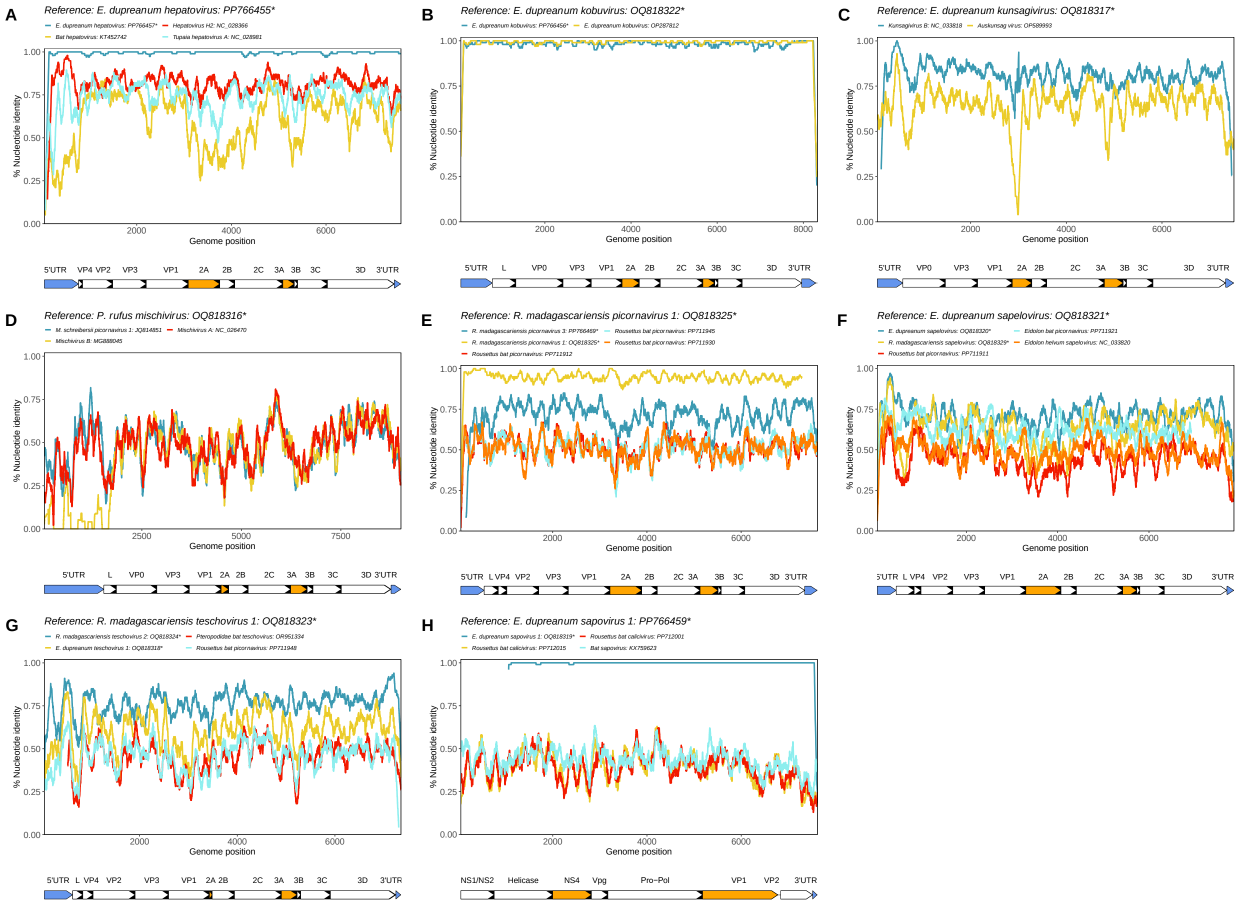

**Supplemental figure 4**: Nucleotide similarity computed in PySimPlot[59] for novel full-length sequences. Similarity analyses with query sequence (A) *E. dupreanum hepatovirus*: accession PP766455, (B) *E. dupreanum kobuvirus*: accession OQ818322, (C) *E. dupreanum kunsagivirus*: accession OQ818317, (D) *P. rufus mischivirus*: accession OQ818316, (E) *R. madagascariensis picornavirus* 1: accession OQ818325, (F) *E. dupreanum sapelovirus*: accession OQ818321, (G) *R. madagascariensis teschovirus* 1: accession OQ818323, and (H) *E. dupreanum sapovirus* 1: accession PP766459 against similar sequences identified from BLAST[55] and other matched novel sequences within the same genus. Asterisks denote novel sequences described in this study. Line color corresponds to different virus sequences, with annotated regions of the genome below each plot. Peptides in orange and corresponding grey shaded areas denote areas of interest for host interactions and immunogenicity, and blue peptides denote 5' and 3' UTRs. Plots were generated with a window size of 200bp and a step size of 20bp.

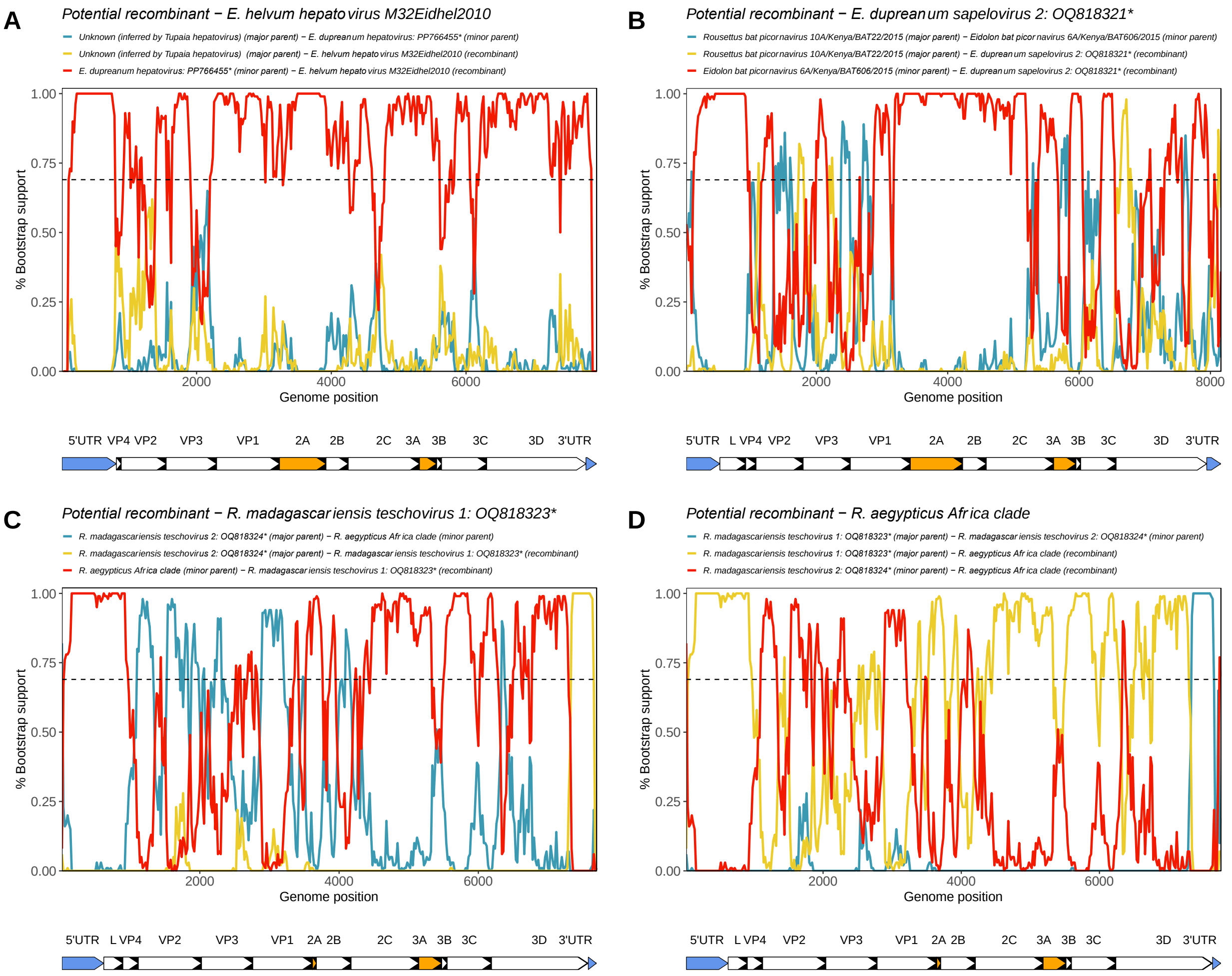


**Supplemental figure 5**: Bootscan plots computed in RDP4[60] for potential recombinant sequences (A) *E. helvum hepatovirus* M32Eidhel2010: accession NC_028366, (B) *E. dupreanum sapelovirus* 2: accession OQ818321, (C) *R. madagascariensis teschovirus* 1: OQ818323, and (D) *R. aegypticus* Africa clade (accessions: PP711948 and PP711934). Line color corresponds to pairwise alignments between the potential recombinant sequence, major parental sequence, and the minor parental sequence. Asterisks denote novel sequences described in this study. Horizontal dashed line refers to a 70% cutoff bootstrap percentage, and grey bars indicate regions identified as significant areas of recombination ($P<0.05$) across at least 5 analyses within RDP4[60] (RDP, GENECONV, Bootscan, Maxchi, Chimaera, and 3Seq). Nucleotide bootscan plots were generated using a window size of 200bp and a step size of 20bp. Genome maps are below each plot, peptides in orange denote areas of interest for host interactions and immunogenicity, and blue peptides denote 5' and 3' UTRs. Full RDP4[60] statistics are in **Supplemental table 6**.
