## Supplemental tables for "*Picornaviridae* and *Caliciviridae* diversity in Madagascar fruit bats is driven by cross-continental genetic exchange"

**Supplemental table 1**: Summary information of phylogenies presented in **Figure 2 and 3**.

| Phylogeny | Figure | Alignment file | # Novel seq | # Reference seq | Region | TaxID references | Overlap length (bp) | Best model |
|---|---|---|---|---|---|---|---|---|
| Summary polymerase | 2A | polymerase_summary_align_trim | 20 | 273 | Polymerase | All taxid listed below | ~2900 | GTR+I+G4 |
| *Cardiovirus* | 2B | cardio_all_align | 1 | 75 | Full genome | 12103, 434308 | ~3600 | TIM2+G4 |
| *Hepatovirus* | 2C | hepato_all_align | 3 | 126 | Full genome | 12091, 1714618 | ~4500 | GTR+I+G4 |
| *Kobuvirus* | 2D | kobu_all_align | 4 | 179 | Full genome | 194960, 655314 | ~1090 | GTR+I+G4 |
| *Kunsagivirus* | 2E | kunsagi_all_align | 1 | 4 | Full genome | 1755589 | ~6700 | TIM2+G4 |
| *Mischivirus* | 2F | mischi_full_align | 1 | 16 | Full genome | 1511778 | ~3200 | GTR+G4 |
| *Unclassified bat picornavirus* | 2G | bat_picorna_all_align | 4 | 35 | Full genome | 2169640, 2788549, 1281456 | ~1700 | GTR+G4 |
| *Sapelovirus* | 2H | sapelo_all_align | 3 | 57 | Full genome | 686982, 1073966 | ~7200 | GTR+G4 |
| *Teschovirus* | 2I | tescho_full_align | 3 | 35 | Full genome | 118139, 2004714 | ~3200 | TIM2+G4 |
| *Sapovirus* | 2J | sapo_all_align | 5 | 217 | Full genome | 95341, 371833 | ~1800 | GTR+I+G4 |

**Supplemental table 2**: Summary table of BLASTx[55] and BLASTn[55] results of novel full and partial-length *Picornaviridae* sequences recovered from mNGS. Bold denotes full-length sequences.

| Accession | Virus | BLASTx host species | BLASTx coverage (%) | BLASTx identity (%) | BLASTx accession | BLASTn host species | BLASTn coverage (%) | BLASTn identity (%) | BLASTn accession |
|---|---|---|---|---|---|---|---|---|---|
| PP766467 | *E. dupreanum cardiovirus* | *Pongo* | 91 | 95.42 | QMI58083.1 | *Rattus norvegicus* | 97 | 83.04 | DQ835185.2 |
| OQ818337 | *E. dupreanum hepatovirus* | *Eidolon helvum* | 87 | 94.75 | YP_009179216.1 | *Eidolon helvum* | 97 | 82.4 | NC_028366.1 |
| PP766455 | *E. dupreanum hepatovirus* | *Eidolon helvum* | 91 | 94.3 | YP_009179216.1 | *Eidolon helvum* | 78 | 99.54 | NC_028366.1 |
| PP766457 | *E. dupreanum hepatovirus* | *Eidolon helvum* | 90 | 94.26 | YP_009179216.1 | *Eidolon helvum* | 97 | 82.16 | NC_028366.1 |
| PP766458 | *E. dupreanum hepatovirus* | *Eidolon helvum* | 99 | 92.68 | YP_009179216.1 | *Eidolon helvum* | 99 | 81.53 | NC_028366.1 |
| **OQ818322** | *E. dupreanum kobuvirus* | *Eidolon dupreanum* | 87 | 100 | WBP49885.1 | *Eidolon dupreanum* | 98 | 99.42 | OP287812.1 |
| PP766449 | *E. dupreanum kobuvirus* | *Eidolon helvum* | 69 | 99.33 | AGL97808.1 | *Eidolon dupreanum* | 100 | 98.99 | OP287812.1 |
| PP766451 | *E. dupreanum kobuvirus* | *Eidolon dupreanum* | 99 | 99.47 | WBP49885.1 | *Eidolon dupreanum* | 99 | 98.48 | OP287812.1 |
| PP766452 | *E. dupreanum kobuvirus* | *Eidolon dupreanum* | 99 | 99.31 | WBP49885.1 | *Eidolon dupreanum* | 100 | 97.88 | OP287812.1 |
| PP766453 | *E. dupreanum kobuvirus* | *Eidolon dupreanum* | 35 | 97.54 | WBP49885.1 | *Eidolon dupreanum* | 99 | 99.12 | OR082796.1 |
| PP766454 | *E. dupreanum kobuvirus* | *Eidolon dupreanum* | 85 | 99.81 | WBP49885.1 | *Eidolon dupreanum* | 100 | 98.45 | OP287812.1 |
| **PP766456** | *E. dupreanum kobuvirus* | *Eidolon dupreanum* | 88 | 99.71 | WBP49885.1 | *Eidolon dupreanum* | 99 | 98.4 | OP287812.1 |
| PP766450 | *E. dupreanum kobuvirus 2* | *Eidolon dupreanum* | 83 | 99.72 | WBP49885.1 | *Eidolon dupreanum* | 85 | 97.91 | OP287812.1 |
| **OQ818317** | *E. dupreanum kunsagivirus* | *Eidolon helvum* | 90 | 91.61 | YP_009345896.1 | *Eidolon helvum* | 95 | 81.93 | NC_033818.1 |
| **OQ818316** | *P. rufus mischivirus* | *Hipposideros gigas* | 81 | 46.51 | YP_009121743.1 | *Suncus murinus* | 10 | 69.71 | OQ716013.1 |
| OQ818325 | *R. madagascariensis picornavirus 1* | *Rousettus aegyptiacus* | 91 | 79.35 | XBH24017.1 | *Rousettus aegyptiacus* | 43 | 73.96 | PP711945.1 |
| **OQ818328** | *R. madagascariensis picornavirus 1* | *Rousettus aegyptiacus* | 91 | 79.7 | XBH24017.1 | *Rousettus aegyptiacus* | 97 | 73.02 | PP711913.1 |
| OQ818346 | *R. madagascariensis picornavirus 2* | *Rousettus aegyptiacus* | 99 | 81.82 | XBH23984.1 | *Bos taurus* | 15 | 81.94 | ON168930.1 |
| **PP766469** | *R. madagascariensis picornavirus 3* | *Rousettus aegyptiacus* | 92 | 79.03 | XBH24017.1 | *Rousettus aegyptiacus* | 51 | 72.67 | PP711909.1 |
| PP766471 | *R. madagascariensis picornavirus 3* | *Rousettus aegyptiacus* | 82 | 82.11 | XBH23984.1 | *Rousettus aegyptiacus* | 99 | 74.6 | PP711913.1 |
| PP766472 | *R. madagascariensis picornavirus 3* | *Rousettus aegyptiacus* | 99 | 74.54 | XBH24017.1 | *Rousettus aegyptiacus* | 38 | 75.11 | PP711928.1 |
| PP766475 | *R. madagascariensis picornavirus 3* | *Rousettus aegyptiacus* | 99 | 78.65 | XBH24017.1 | *Rousettus aegyptiacus* | 99 | 73.99 | PP711909.1 |
| **OQ818320** | *E. dupreanum sapelovirus 1* | *Eidolon helvum* | 96 | 98.95 | YP_009345901.1 | *Eidolon helvum* | 99 | 94.37 | KX644938.1 |
| OQ818342 | *E. dupreanum sapelovirus 1* | *Eidolon helvum* | 99 | 98.47 | YP_009345901.1 | *Eidolon helvum* | 99 | 94.73 | KX644938.1 |
| OQ818343 | *E. dupreanum sapelovirus 1* | *Eidolon helvum* | 99 | 98.67 | YP_009345901.1 | *Eidolon helvum* | 100 | 92.28 | KX644938.1 |
| OQ818344 | *E. dupreanum sapelovirus 1* | *Eidolon helvum* | 99 | 99.15 | YP_009345901.1 | *Eidolon helvum* | 99 | 94.57 | KX644938.1 |
| PP766465 | *E. dupreanum sapelovirus 1* | *Eidolon helvum* | 99 | 99.19 | YP_009345901.1 | *Eidolon helvum* | 100 | 94.24 | KX644938.1 |
| PP766466 | *E. dupreanum sapelovirus 1* | *Eidolon helvum* | 99 | 98.91 | YP_009345901.1 | *Eidolon helvum* | 100 | 92.83 | KX644938.1 |
| **OQ818321** | *E. dupreanum sapelovirus 2* | *Eidolon helvum* | 83 | 97.81 | XBH23993.1 | *Eidolon helvum* | 87 | 87.42 | PP711921.1 |
| PP766462 | *E. dupreanum sapelovirus 2* | *Eidolon helvum* | 99 | 97.03 | XBH23993.1 | *Eidolon helvum* | 100 | 86.55 | PP711921.1 |
| PP766463 | *E. dupreanum sapelovirus 2* | *Eidolon helvum* | 99 | 98.31 | XBH23993.1 | *Eidolon helvum* | 100 | 88.18 | PP711921.1 |
| PP766464 | *E. dupreanum sapelovirus 2* | *Eidolon helvum* | 99 | 98.45 | XBH23993.1 | *Eidolon helvum* | 100 | 88.58 | PP711921.1 |
| **OQ818329** | *R. madagascariensis sapelovirus 1* | *Rousettus aegyptiacus* | 94 | 89.68 | XBH23983.1 | *Rousettus aegyptiacus* | 97 | 79.04 | PP711911.1 |
| **OQ818318** | *E. dupreanum teschovirus 1* | *Rousettus aegyptiacus* | 90 | 73.14 | XBH24020.1 | *Rousettus aegyptiacus* | 32 | 72.67 | PP711948.1 |
| **OQ818323** | *R. madagascariensis teschovirus 1* | *Rousettus aegyptiacus* | 87 | 92.16 | XBH24006.1 | *Rousettus aegyptiacus* | 94 | 82.69 | PP711934.1 |
| **OQ818324** | *R. madagascariensis teschovirus 2* | *Rousettus aegyptiacus* | 88 | 92.25 | XBH24006.1 | *Rousettus aegyptiacus* | 94 | 79.46 | PP711934.1 |

**Supplemental table 3**: Summary table of BLASTx[55] and BLASTn[55] results of novel full and partial-length *Caliciviridae* sequences recovered from mNGS. Bold denotes full-length sequences.

| Accession | Virus | BLASTx host species | BLASTx coverage (%) | BLASTx identity (%) | BLASTx accession | BLASTn host species | BLASTn coverage (%) | BLASTn identity (%) | BLASTn accession |
|---|---|---|---|---|---|---|---|---|---|
| OQ818319 | *E. dupreanum sapovirus 1* | *Eidolon helvum* | 90 | 67.18 | AQQ78883.1 | *Homo sapiens* | 1 | 79.82 | LC504397.1 |
| **PP766459** | *E. dupreanum sapovirus 1* | *Eidolon helvum* | 77 | 99.9 | AQQ78883.1 | *Homo sapiens* | 1 | 79.82 | LC504397.1 |
| OQ818340 | *E. dupreanum sapovirus 2* | *Eidolon helvum* | 99 | 67.97 | AQQ78883.1 | *Eidolon helvum* | 63 | 71.4 | KX759619.1 |
| PP766461 | *E. dupreanum sapovirus 3* | *Eidolon helvum* | 96 | 90.13 | AQQ78883.1 | *Eidolon helvum* | 92 | 80.7 | KX759623.1 |
| PP766460 | *E. dupreanum sapovirus 4* | *Eidolon helvum* | 70 | 75.26 | AQQ78883.1 | No result | No result | No result | No result |
| OQ818345 | *R. madagascariensis sapovirus 1* | *Rousettus aegyptiacus* | 99 | 68.19 | XBH24168.1 | No result | No result | No result | No result |
| OQ818347 | *R. madagascariensis sapovirus 2* | *Rousettus aegyptiacus* | 99 | 89.63 | XBH24156.1 | *Rousettus aegypticus* | 100 | 77.94 | PP712001.1 |
| PP766470 | *R. madagascariensis sapovirus 2* | *Rousettus aegyptiacus* | 99 | 90.86 | XBH24177.1 | *Rousettus aegypticus* | 99 | 78.51 | PP712015.1 |
| PP766473 | *R. madagascariensis sapovirus 2* | *Rousettus aegyptiacus* | 99 | 90.59 | XBH24177.1 | *Rousettus aegypticus* | 97 | 77.28 | PP712015.1 |
| PP766474 | *R. madagascariensis sapovirus 2* | *Rousettus aegyptiacus* | 100 | 94.55 | XBH24177.1 | *Rousettus aegypticus* | 68 | 71.74 | PP712026.1 |
| PP766476 | *R. madagascariensis sapovirus 2* | *Rousettus aegyptiacus* | 99 | 86.05 | XBH24177.1 | *Eidolon helvum* | 19 | 73.71 | KX759623.1 |
| PP766477 | *R. madagascariensis sapovirus 2* | *Rousettus aegyptiacus* | 53 | 81.32 | XBH24178.1 | *Rousettus aegypticus* | 72 | 76.22 | PP712001.1 |
| OQ818348 | *R. madagascariensis sapovirus 3* | *Rousettus aegyptiacus* | 99 | 84.12 | XBH24163.1 | *Rousettus aegypticus* | 94 | 75 | PP712006.1 |
| PP766468 | *R. madagascariensis sapovirus 3* | *Rousettus aegyptiacus* | 100 | 85 | XBH24156.1 | *Rousettus aegypticus* | 87 | 75.67 | PP712033.1 |

**Supplemental table 4**: Peptide cleavage sites for full and partial-length *Picornaviridae* sequences described in this study. Bold denotes full-length sequences.

| Accession | Virus | L/VP4 or VP0 | VP4/VP2 | VP2 or VP0/VP3 | VP3/VP1 | VP1/2A | 2A/2B | 2B/2C | 2C/3A | 3A/3B | 3B/3C | 3C/3D |
|---|---|---|---|---|---|---|---|---|---|---|---|---|
| PP766467 | *E. dupreanum cardiovirus* | - | - | - | - | - | - | qqG/Spl | vaQ/Apv | qeQ/Gpy | diQ/Gpn | epQ/Gal |
| OQ818337 | *E. dupreanum hepatovirus* | - | tlA/Die | mtQ/Mmr | ttQ/Agd | kfE/Eel | ssE/Ase | kaE/Sld | wsQ/Gfs | ltT/Gvy | dsQ/Svv | - |
| PP766455 | *E. dupreanum hepatovirus* | - | - | - | - | - | - | - | lfQ/Ggv | gsQ/Gpy | - | - |
| PP766457 | *E. dupreanum hepatovirus* | kpN/Fah | tlK/Spt | saQ/Gfp | - | - | - | - | - | - | - | - |
| PP766458 | *E. dupreanum hepatovirus* | - | - | - | - | - | - | - | lfQ/Ggv | gsQ/Gpy | enQ/Gpd | giQ/Gvi |
| **OQ818322** | *E. dupreanum kobuvirus* | qrQ/Gns | - | akQ/Hwk | ssQ/Ags | vkQ/Gat | rrQ/Gll | qtQ/Glr | krQ/Grv | qpQ/Aay | qrG/Gms | qqQ/Sli |
| PP766449 | *E. dupreanum kobuvirus* | sgQ/Gty | llA/Qps | wrG/Sit | svQ/Dsn | vqQ/Apk | npG/Pai | nkQ/Gkl | skQ/Apt | hnQ/Gpy | elQ/Apg | akQ/Gki |
| PP766451 | *E. dupreanum kobuvirus* | kpM/Yah | alK/Spt | aaQ/Gip | ffQ/Glg | yyQ/Eqa | veQ/Gll | eeQ/Gpt | lfQ/Gpg | gqQ/Gpy | evQ/Gpd | nfQ/Gsi |
| PP766452 | *E. dupreanum kobuvirus* | qrQ/Gns | - | akQ/Hwk | ssQ/Ags | vkQ/Gat | rrQ/Gll | qtQ/Glr | krQ/Grv | qpQ/Aay | qrG/Gms | qqQ/Sli |
| PP766453 | *E. dupreanum kobuvirus* | tqC/Gva | alK/Spt | srQ/Gvp | yfQ/Glg | isQ/Tpi | meQ/Gpm | lpQ/Gvs | lfQ/Gpv | ssQ/Gpy | vtQ/Gpd | leQ/Gei |
| PP766454 | *E. dupreanum kobuvirus* | rqC/Gva | alK/Spt | mrQ/Gvp | yfQ/Glg | igQ/Tpi | meQ/Gpv | vqQ/Gvs | lfQ/Gpp | snQ/Gpy | vtQ/Gpd | leQ/Gqi |
| PP766456 | *E. dupreanum kobuvirus* | hkQ/Gvg | llA/Dpk | ytE/Gpa | gtQ/Gpe | vqQ/Gpr | npG/Pai | nkQ/Gkl | skQ/Apq | hnQ/Gpy | elQ/Apg | tkQ/Gki |
| PP766450 | *E. dupreanum kobuvirus 2* | rqC/Gva | alK/Spt | mrQ/Gvp | yfQ/Glg | igQ/Tpi | meQ/Gpv | vqQ/Gvs | lfQ/Gpp | snQ/Gpy | vtQ/Gpd | leQ/Gqi |
| **OQ818317** | *E. dupreanum kunsagivirus* | - | - | rrQ/Ifs | lyH/App | aaQ/Gpr | iyG/Pti | ipQ/Gpf | faQ/Sps | psQ/Gpy | dpQ/Gpw | raQ/Gqi |
| **OQ818316** | *P. rufus mischivirus* | mfQ/Gag | - | LRQ/Pkl | slQ/Tsl | ttE/Gsp | npG/Pet | klQ/Pll | eiQ/Tke | enQ/Apy | neQ/Kpr | lsE/Gvd |
| OQ818325 | *R. madagascariensis picornavirus 1* | rqC/Gva | alK/Spt | mrQ/Gvp | yfQ/Glg | igQ/Tpi | meQ/Gpv | vqQ/Gvs | lfQ/Gpp | snQ/Gpy | vtQ/Gpd | leQ/Gqi |
| **OQ818328** | *R. madagascariensis picornavirus 1* | rqC/Gva | alK/Spt | mrQ/Gvp | yfQ/Glg | igQ/Tpi | meQ/Gpv | vqQ/Gvs | lfQ/Gpp | snQ/Gpy | vtQ/Gpd | leQ/Gqi |
| OQ818346 | *R. madagascariensis picornavirus 2* | - | tlA/Die | mtQ/Mmr | ttQ/Agd | kfE/Eel | ssE/Ase | kaE/Sld | wsQ/Gfs | ltT/Gvy | dsQ/Svv | - |
| **PP766469** | *R. madagascariensis picornavirus 3* | kpM/Yah | alK/Spt | aaQ/Gip | ffQ/Glg | yyQ/Eqa | veQ/Gll | eeQ/Gpt | lfQ/Gpg | gqQ/Gpy | evQ/Gpd | nfQ/Gsi |
| PP766471 | *R. madagascariensis picornavirus 3* | - | - | - | - | - | - | - | lfQ/Ggv | gsQ/Gpy | - | - |
| PP766472 | *R. madagascariensis picornavirus 3* | kpN/Fah | tlK/Spt | saQ/Gfp | - | - | - | - | - | - | - | - |
| PP766475 | *R. madagascariensis picornavirus 3* | - | - | - | - | - | - | - | lfQ/Ggv | gsQ/Gpy | enQ/Gpd | giQ/Gvi |
| **OQ818320** | *E. dupreanum sapelovirus 1* | kpN/Fah | tlK/Spt | saQ/Gfp | dlQ/Gfi | irL/Gpp | vyQ/Gik | eeQ/Gft | lfQ/Ggv | gsQ/Gpy | evQ/Gpd | giQ/Gvi |
| OQ818342 | *E. dupreanum sapelovirus 1* | hkQ/Gvg | llA/Dpk | ytE/Gpa | gtQ/Gpe | vqQ/Gpr | npG/Pai | nkQ/Gkl | skQ/Apq | hnQ/Gpy | elQ/Apg | tkQ/Gki |
| OQ818343 | *E. dupreanum sapelovirus 1* | sgQ/Gty | llA/Qps | wrG/Sit | svQ/Dsn | vqQ/Apk | npG/Pai | nkQ/Gkl | skQ/Apt | hnQ/Gpy | elQ/Apg | akQ/Gki |
| OQ818344 | *E. dupreanum sapelovirus 1* | rqC/Gva | alK/Spt | mrQ/Gvp | yfQ/Glg | igQ/Tpi | meQ/Gpv | vqQ/Gvs | lfQ/Gpp | snQ/Gpy | vtQ/Gpd | leQ/Gqi |
| PP766465 | *E. dupreanum sapelovirus 1* | rqC/Gva | alK/Spt | mrQ/Gvp | yfQ/Glg | igQ/Tpi | meQ/Gpv | vqQ/Gvs | lfQ/Gpp | snQ/Gpy | vtQ/Gpd | leQ/Gqi |
| PP766466 | *E. dupreanum sapelovirus 1* | - | tlA/Die | mtQ/Mmr | ttQ/Agd | kfE/Eel | ssE/Ase | kaE/Sld | wsQ/Gfs | ltT/Gvy | dsQ/Svv | - |
| **OQ818321** | *E. dupreanum sapelovirus 2* | kpN/Iah | alK/Spt | stQ/Gip | evQ/Gfv | irL/Gpp | eiQ/Gik | eeQ/Glv | lfQ/Gpi | gsQ/Gpy | evQ/Gpd | sfQ/Gki |
| PP766462 | *E. dupreanum sapelovirus 2* | kpM/Yah | alK/Spt | aaQ/Gip | ffQ/Glg | yyQ/Eqa | veQ/Gll | eeQ/Gpt | lfQ/Gpg | gqQ/Gpy | evQ/Gpd | nfQ/Gsi |
| PP766463 | *E. dupreanum sapelovirus 2* | qrQ/Gns | - | akQ/Hwk | ssQ/Ags | vkQ/Gat | rrQ/Gll | qtQ/Glr | krQ/Grv | qpQ/Aay | qrG/Gms | qqQ/Sli |
| PP766464 | *E. dupreanum sapelovirus 2* | tqC/Gva | alK/Spt | srQ/Gvp | yfQ/Glg | isQ/Tpi | meQ/Gpm | lpQ/Gvs | lfQ/Gpv | ssQ/Gpy | vtQ/Gpd | leQ/Gei |
| **OQ818329** | *R. madagascariensis sapelovirus 1* | qrQ/Gns | - | akQ/Hwk | ssQ/Ags | vkQ/Gat | rrQ/Gll | qtQ/Glr | krQ/Grv | qpQ/Aay | qrG/Gms | qqQ/Sli |
| **OQ818318** | *E. dupreanum teschovirus 1* | hkQ/Gag | lsS/Gln | alQ/Gpi | siQ/Gnt | tkQ/Gat | npG/Ppv | kkQ/Gll | tkQ/Apk | keQ/Say | qlQ/Apg | ekQ/Gki |
| **OQ818323** | *R. madagascariensis teschovirus 1* | kpN/Fah | tlK/Spt | saQ/Gfp | dlQ/Gfi | irL/Gpp | vyQ/Gik | eeQ/Gft | lfQ/Ggv | gsQ/Gpy | evQ/Gpd | giQ/Gvi |
| **OQ818324** | *R. madagascariensis teschovirus 2* | kpN/Iah | alK/Spt | stQ/Gip | evQ/Gfv | irL/Gpp | eiQ/Gik | eeQ/Glv | lfQ/Gpi | gsQ/Gpy | evQ/Gpd | sfQ/Gki |

**Supplemental table 5**: Peptide cleavage sites for full and partial-length *Caliciviridae* sequences described in this study. Bold denotes full-length sequences.

| Accession | Virus | NS1 and NS2/Helicase | Helicase/NS4 | NS4/Vpg | Vpg/Pro-Pol |
|---|---|---|---|---|---|
| OQ818319 | *E. dupreanum sapovirus 1* | - | eaQ/Agk | giE/Akg | esQ/Ags |
| **PP766459** | *E. dupreanum sapovirus 1* | qpQ/Aia | eaQ/Agk | giE/Akg | esQ/Ags |
| OQ818340 | *E. dupreanum sapovirus 2* | - | eaQ/Agk | gvE/Akg | esQ/Ant |
| PP766461 | *E. dupreanum sapovirus 3* | - | - | - | - |
| PP766460 | *E. dupreanum sapovirus 4* | - | - | - | - |
| OQ818345 | *R. madagascariensis sapovirus 1* | - | - | - | - |
| OQ818347 | *R. madagascariensis sapovirus 2* | - | - | gkK/Gkt | epE/Sgd |
| PP766470 | *R. madagascariensis sapovirus 2* | - | - | tdE/Akg | epE/San |
| PP766473 | *R. madagascariensis sapovirus 2* | - | - | tdE/Akg | epE/San |
| PP766474 | *R. madagascariensis sapovirus 2* | - | eaQ/Apn | - | - |
| PP766476 | *R. madagascariensis sapovirus 2* | - | - | - | - |
| PP766477 | *R. madagascariensis sapovirus 2* | - | - | - | - |
| OQ818348 | *R. madagascariensis sapovirus 3* | - | fpQ/Ssd | eeE/Akg | idE/Gps |
| PP766468 | *R. madagascariensis sapovirus 3* | - | eaQ/Sgn | - | - |

**Supplemental table 6**: RDP4[60] output of comparisons involving novel sequences as either the recombinant sequence or a major/minor parental sequence. Unknown followed by an accession number indicates that RDP4[60] used that named sequence to infer an unknown parental sequence. NS denotes non-significance in that analysis. Recombination events are accepted if at least 4 tests are significant. Recombinant sequences with highest support are highlighted in red (5-6 significant tests). Novel sequences identified in this study are in bold. Particular lineages are consensus sequences of aligned phylogenetic clade members and include: *Rhinolophus bat picornavirus* BtSY4 clade (accessions OP963617 and PP746000), Shanbavirus A clade (accessions KJ641687, KJ641690, KJ641699, and KJ641693), *R. aegypticus* African picornavirus clade (accessions PP711912, PP711945, PP711913, PP711930, PP711928, and PP711909), *H. larvatus* Chinese hepatovirus clade (accessions OR951274, OR951275, OR867086, OR951271), *Rattus tanezumi* Chinese kobuvirus clade (accessions OM069746 and OM069755), *Rhinolophus* Chinese kobuvirus clade (accessions OR951245 and OR951365), Rodent Chinese kunsagivirus clade (accessions OQ716009 and OQ716008), *Eonycteris* Chinese sapelovirus clade (accessions OR951327, OR951326, OR951325, and OR951332), *E. helvum* Cameroonian sapovirus clade (accessions KX759619, KX759623, KX759621, NC_033776, and KX759622), *R. aegypticus* Kenyan sapovirus clade (accessions PP712001, PP712004, PP712008, and PP712015), *R. aegypticus* east African teschovirus clade (accessions PP711948 and PP711934), and *Rousettus* Chinese teschovirus clade (accessions OR951335, OR951333, and OR951334).

| Alignment file | Genus | Breakpoint Begin | Breakpoint End | Recombinant Sequence | Major Parental Sequence | Minor Parental Sequence | RDP | GENECONV | Bootscan | Maxchi | Chimaera | 3Seq | Number significant tests | Accept Recombination Event? | Corresponding Figure |
|---|---|---|---|---|---|---|---|---|---|---|---|---|---|---|---|
| batpicorna_rdp_align_nt_clade | *Bat picornavirus* | 117 | 184 | rhinolophus_BtSY4 | **Unknown (OQ818328)** | PP711912 | 7.56E-03 | NS | NS | 8.86E-03 | 1.28E-03 | NS | 2 | no | |
| batpicorna_rdp_align_nt_clade | *Bat picornavirus* | | | | **Unknown(OQ818325)** | | | | | | | | | | |
| batpicorna_rdp_align_nt_clade | *Bat picornavirus* | 6885 | 7153 | rhinolophus_BtSY4 | **OQ818325** | **Unknown (OQ818328)** | NS | NS | 9.66E-03 | NS | NS | NS | 1 | no | |
| batpicorna_rdp_align_nt_clade | *Bat picornavirus* | 3238 | 3452 | **PP766469** | PP711912 | **Unknown (OQ818325)** | 1.70E-02 | NS | 2.07E-02 | NS | NS | NS | 2 | no | |
| batpicorna_rdp_align_nt_clade | *Bat picornavirus* | 962 | 1311 | **PP766469** | rhinolophus_BtSY4 | **OQ818325** | NS | NS | 1.27E-02 | 3.42E-04 | 1.71E-02 | NS | 3 | no | |
| batpicorna_rdp_align_nt_clade | *Bat picornavirus* | | | | | **OQ818328** | | | | | | | | | |
| batpicorna_rdp_align_nt_clade | *Bat picornavirus* | 1548 | 1709 | rhinolophus_BtSY4 | shanbavirus_a | **PP766469** | 1.52E-02 | NS | NS | NS | NS | NS | 1 | no | |
| batpicorna_rdp_align_nt | *Bat picornavirus* | 844 | 1148 | OQ363775.1 | Unknown (PP746000) | **PP766469** | 2.21E-04 | NS | NS | NS | 3.30E-03 | NS | 2 | no | |
| batpicorna_rdp_align_nt | *Bat picornavirus* | | | HQ595345.1 | | | | | | | | | | | |
| batpicorna_rdp_align_nt | *Bat picornavirus* | | | NC_015934.1 | | | | | | | | | | | |
| batpicorna_rdp_align_nt | *Bat picornavirus* | | | KJ641693.1 | | | | | | | | | | | |
| batpicorna_rdp_align_nt | *Bat picornavirus* | | | OQ363764.1 | | | | | | | | | | | |
| batpicorna_rdp_align_nt | *Bat picornavirus* | 1365 | 1464 | PP711930 | **OQ818328** | KJ641687.1 | 6.84E-04 | NS | NS | 2.84E-03 | 1.74E-04 | NS | 3 | no | |
| batpicorna_rdp_align_nt | *Bat picornavirus* | | | PP711909 | **OQ818325** | KJ641690.1 | | | | | | | | | |
| batpicorna_rdp_align_nt | *Bat picornavirus* | 3218 | 4049 | HQ595345.1 | **OQ818328** | **Unknown (PP766469)** | 6.01E-04 | 3.60E-03 | NS | 7.20E-04 | 4.10E-02 | NS | 4 | yes | |
| batpicorna_rdp_align_nt | *Bat picornavirus* | | | NC_015934.1 | | | | | | | | | | | |
| batpicorna_rdp_align_nt | *Bat picornavirus* | | | OQ363775.1 | | | | | | | | | | | |
| batpicorna_rdp_align_nt | *Bat picornavirus* | 5057 | 5220 | **PP766469** | **OQ818328** | Unknown (PP746000) | 7.56E-04 | NS | NS | NS | NS | NS | 1 | no | |
| batpicorna_rdp_align_nt | *Bat picornavirus* | 1588 | 1689 | PP746000 | OP963617.1 | **PP766469** | 1.97E-03 | NS | NS | NS | NS | NS | 1 | no | |
| batpicorna_rdp_align_nt | *Bat picornavirus* | 330 | 554 | **OQ818328** | **PP766469** | OP963617.1 | 3.55E-03 | NS | NS | NS | NS | 0.01616447 | 2 | no | |
| batpicorna_rdp_align_nt | *Bat picornavirus* | | | **OQ818325** | | | | | | | | | | | |
| batpicorna_rdp_align_nt | *Bat picornavirus* | 4688 | 4860 | **OQ818328** | **PP766469** | PP745912 | 6.88E-03 | NS | NS | NS | NS | 4.62E-02 | 2 | no | |
| batpicorna_rdp_align_nt | *Bat picornavirus* | | | **OQ818325** | | | | | | | | | | | |
| batpicorna_rdp_align_nt | *Bat picornavirus* | 6704 | 6859 | PP745912 | Unknown (HQ595345.1) | **OQ818328** | 5.38E-03 | NS | 3.43E-02 | NS | NS | NS | 2 | no | |
| batpicorna_rdp_align_nt | *Bat picornavirus* | | | | Unknown(NC_015934.1) | | | | | | | | | | |
| batpicorna_rdp_align_nt | *Bat picornavirus* | 7387 | 34 | PP711912 | PP711930 | **Unknown (OQ818328)** | 1.35E-02 | NS | 0.01884588 | 1.01E-02 | 5.34E-03 | NS | 4 | yes | |
| batpicorna_rdp_align_nt | *Bat picornavirus* | 117 | 201 | OP963617.1 | **OQ818328** | Unknown (PP711945) | 0.0192717 | NS | NS | NS | NS | NS | 1 | no | |
| batpicorna_rdp_align_nt | *Bat picornavirus* | 3262 | 3281 | **OQ818325** | PP711928 | Unknown (OQ363764.1) | 2.23E-02 | NS | NS | NS | NS | NS | 1 | no | |
| batpicorna_rdp_align_nt | *Bat picornavirus* | | | **OQ818328** | | | | | | | | | | | |
| batpicorna_rdp_align_nt | *Bat picornavirus* | 5268 | 5303 | **OQ818328** | PP711928 | HQ595345.1 | 2.36E-02 | NS | NS | NS | NS | NS | 1 | no | |
| batpicorna_rdp_align_nt | *Bat picornavirus* | | | **OQ818325** | | NC_015934.1 | | | | | | | | | |
| batpicorna_rdp_align_nt | *Bat picornavirus* | 4854 | 4882 | PP711928 | **OQ818328** | KJ641693.1 | 2.24E-02 | NS | NS | NS | NS | NS | 1 | no | |
| batpicorna_rdp_align_nt | *Bat picornavirus* | | | | **OQ818325** | | | | | | | | | | |
| batpicorna_rdp_align_nt | *Bat picornavirus* | 6448 | 6524 | OQ363764.1 | **PP766469** | KJ641690.1 | 2.68E-02 | NS | NS | NS | NS | NS | 1 | no | |
| batpicorna_rdp_align_nt | *Bat picornavirus* | | | HQ595345.1 | | KJ641687.1 | | | | | | | | | |
| batpicorna_rdp_align_nt | *Bat picornavirus* | | | NC_015934.1 | | KJ641699.1 | | | | | | | | | |
| batpicorna_rdp_align_nt | *Bat picornavirus* | | | OQ363775.1 | | | | | | | | | | | |
| batpicorna_rdp_align_nt | *Bat picornavirus* | 3859 | 3987 | PP746000 | **Unknown (OQ818328)** | KJ641690.1 | 2.79E-02 | NS | NS | NS | NS | NS | 1 | no | |
| batpicorna_rdp_align_nt | *Bat picornavirus* | 2654 | 3026 | PP711946 | Unknown (KJ641690.1) | **PP766469** | 4.87E-02 | NS | NS | 3.90E-02 | NS | NS | 2 | no | |
| hepato_rdp_align_nt_clade | *Hepatovirus* | 7198 | 560 | NC_028366 | Unknown (NC_028981) | **PP766455** | 1.83E-04 | 1.46E-06 | 1.58E-08 | 6.59E-06 | 1.51E-02 | NS | 5 | yes | Sfig. 5A |
| hepato_rdp_align_nt_clade | *Hepatovirus* | | | | | **PP766457** | | | | | | | | | |
| hepato_rdp_align_nt_clade | *Hepatovirus* | 577 | 7197 | NC_028366 | **PP766457** | H_larvatus_china | NS | NS | 5.76E-04 | 6.73E-03 | 7.99E-03 | NS | 3 | no | |
| hepato_rdp_align_nt_clade | *Hepatovirus* | | | | **PP766455** | | | | | | | | | | |
| hepato_rdp_align_nt_clade | *Hepatovirus* | 1745 | 1968 | **PP766457** | NC_028366 | KT452729 | 6.03E-03 | NS | NS | NS | 4.27E-02 | 4.02E-02 | 3 | no | |
| hepato_rdp_align_nt_clade | *Hepatovirus* | | | **PP766455** | | | | | | | | | | | |
| hepato_rdp_align_nt_clade | *Hepatovirus* | 1012 | 1156 | NC_038313 | Unknown (H_larvatus_china) | **PP766455** | 1.52E-02 | 2.22E-02 | NS | 9.13E-03 | NS | NS | 3 | no | |
| hepato_rdp_align_nt_clade | *Hepatovirus* | | | | | **PP766457** | | | | | | | | | |
| hepato_rdp_align_nt | *Hepatovirus* | 7224 | 560 | NC_028366 | Unknown (NC_028981) | **PP766455** | 1.50E-03 | 3.63E-05 | 1.92E-07 | 1.96E-04 | 1.36E-04 | NS | 5 | yes | |
| hepato_rdp_align_nt | *Hepatovirus* | | | | | **PP766457** | | | | | | | | | |
| hepato_rdp_align_nt | *Hepatovirus* | 6939 | 640 | OR867086 | **Unknown (PP766455)** | OR951271 | NS | 5.02E-03 | 7.66E-05 | 8.91E-05 | NS | NS | 3 | no | |
| hepato_rdp_align_nt | *Hepatovirus* | | | OR951275 | | | | | | | | | | | |
| hepato_rdp_align_nt | *Hepatovirus* | 3326 | 4084 | NC_038313 | OR951275 | **Unknown (PP766457)** | 5.41E-03 | NS | NS | 2.66E-02 | NS | NS | 2 | no | |
| hepato_rdp_align_nt | *Hepatovirus* | | | | | **Unknown(PP766455)** | | | | | | | | | |
| hepato_rdp_align_nt | *Hepatovirus* | 1583 | 1920 | NC_028366 | **PP766455** | Unknown (KT452729) | 1.09E-02 | NS | NS | NS | NS | NS | 1 | no | |
| hepato_rdp_align_nt | *Hepatovirus* | | | | **PP766457** | | | | | | | | | | |
| kobu_rdp_align_nt_clade | *Kobuvirus* | 7921 | 8347 | **OQ818322** | OP287812.1 | Unknown (NC_034971.1) | NS | NS | 3.08E-04 | NS | NS | NS | 1 | no | |
| kobu_rdp_align_nt_clade | *Kobuvirus* | | | | NC_034971.1 | Unknown(OP287812.1) | | | | | | | | | |
| kobu_rdp_align_nt_clade | *Kobuvirus* | 1482 | 1817 | NC_034971.1 | rattus_tanezumi_china | **Unknown (PP766456)** | NS | NS | 8.99E-04 | NS | NS | 4.45E-03 | 2 | no | |
| kobu_rdp_align_nt_clade | *Kobuvirus* | | | | MN116647 | Unknown(OP287812.1) | | | | | | | | | |
| kobu_rdp_align_nt_clade | *Kobuvirus* | | | | | **Unknown(OQ818322)** | | | | | | | | | |
| kobu_rdp_align_nt_clade | *Kobuvirus* | 3155 | 3399 | NC_034971.1 | rattus_tanezumi_china | **Unknown (PP766456)** | 6.57E-04 | NS | NS | NS | NS | 1.12E-02 | 2 | no | |
| kobu_rdp_align_nt_clade | *Kobuvirus* | | | | | Unknown(OP287812.1) | | | | | | | | | |
| kobu_rdp_align_nt_clade | *Kobuvirus* | | | | | **Unknown(OQ818322)** | | | | | | | | | |
| kobu_rdp_align_nt_clade | *Kobuvirus* | 4286 | 6667 | rattus_tanezumi_china | rhinolophus_china | **PP766456** | 3.28E-03 | NS | NS | NS | NS | NS | 1 | no | |
| kobu_rdp_align_nt_clade | *Kobuvirus* | | | MN116647 | | | | | | | | | | | |
| kobu_rdp_align_nt_clade | *Kobuvirus* | | | NC_034971.1 | | | | | | | | | | | |
| kobu_rdp_align_nt_clade | *Kobuvirus* | | | | Unknown(MN116647) | OP287812.1 | | | | | | | | | |
| kobu_rdp_align_nt_clade | *Kobuvirus* | | | | Unknown(rattus_tanezumi_china) | **PP766456** | | | | | | | | | |

| | | | | | | | | | | | | | | | |
|---|---|---|---|---|---|---|---|---|---|---|---|---|---|---|---|
| kobu_rdp_align_nt_clade | *Kobuvirus* | 2686 | 2864 | NC_034971.1 | rattus_tanezumi_china | **OQ818322** | 2.10E-02 | NS | NS | NS | NS | NS | 2 | no | |
| kobu_rdp_align_nt | *Kobuvirus* | 3107 | 3399 | NC_034971.1 | OR951245.1 | **Unknown (PP766456)** | 1.56E-04 | NS | 5.25E-03 | NS | NS | 3.54E-02 | 3 | no | |
| kobu_rdp_align_nt | *Kobuvirus* | | | | | Unknown(OP287812.1) | | | | | | | | | |
| kobu_rdp_align_nt | *Kobuvirus* | | | | | **Unknown(OQ818322)** | | | | | | | | | |
| kobu_rdp_align_nt | *Kobuvirus* | 1482 | 1817 | NC_034971.1 | OM069746 | **Unknown (OQ818322)** | 4.98E-04 | NS | 5.14E-03 | NS | NS | 8.45E-03 | 3 | no | |
| kobu_rdp_align_nt | *Kobuvirus* | | | | MN116647 | Unknown(OP287812.1) | | | | | | | | | |
| kobu_rdp_align_nt | *Kobuvirus* | | | | OM069755 | **Unknown(PP766456)** | | | | | | | | | |
| kobu_rdp_align_nt | *Kobuvirus* | 7809 | 8216 | NC_034971.1 | **OQ818322** | Unknown (OP287812.1) | NS | NS | 7.69E-04 | NS | NS | NS | 1 | no | |
| kobu_rdp_align_nt | *Kobuvirus* | 3894 | 4511 | OR951365.1 | **Unknown (PP766456)** | NC_034971.1 | NS | NS | 5.71E-03 | 5.00E-03 | 0.00376144 | NS | 3 | no | |
| kobu_rdp_align_nt | *Kobuvirus* | | | | Unknown(OP287812.1) | | | | | | | | | | |
| kobu_rdp_align_nt | *Kobuvirus* | | | | **Unknown(OQ818322)** | | | | | | | | | | |
| kobu_rdp_align_nt | *Kobuvirus* | 3872 | 5081 | OR951245.1 | **Unknown (OQ818322)** | MN116647 | 7.54E-03 | NS | NS | 1.65E-02 | 8.93E-04 | NS | 3 | no | |
| kobu_rdp_align_nt | *Kobuvirus* | | | OR951365.1 | **Unknown(PP766456)** | | | | | | | | | | |
| kobu_rdp_align_nt | *Kobuvirus* | 7147 | 7253 | MN116647 | OR951245.1 | OP287812.1 | 3.51E-02 | 2.43E-03 | 1.02E-02 | NS | NS | NS | 3 | no | |
| kobu_rdp_align_nt | *Kobuvirus* | | | | | **OQ818322** | | | | | | | | | |
| kobu_rdp_align_nt | *Kobuvirus* | 6619 | 6724 | NC_034971.1 | OM069746 | **PP766456** | 0.01107833 | NS | NS | 7.54E-03 | 9.86E-03 | NS | 3 | no | |
| kobu_rdp_align_nt | *Kobuvirus* | 2686 | 2864 | NC_034971.1 | OM069755 | **OQ818322** | 2.26E-02 | NS | 3.23E-02 | NS | NS | NS | 2 | no | |
| kobu_rdp_align_nt | *Kobuvirus* | 7905 | 8156 | OP287812.1 | OR951245.1 | Unknown (MN116647) | NS | 1.57E-03 | 4.74E-02 | 0.02374371 | NS | NS | 3 | no | |
| kobu_rdp_align_nt | *Kobuvirus* | | | **OQ818322** | MN116647 | Unknown(OR951245.1) | | | | | | | | | |
| kobu_rdp_align_nt | *Kobuvirus* | | | **PP766456** | | | | | | | | | | | |
| kobu_rdp_align_nt | *Kobuvirus* | 5085 | 5593 | OR951245.1 | **Unknown (PP766456)** | OM069746 | 3.60E-02 | 1.52E-02 | NS | NS | 2.83E-02 | NS | 3 | no | |
| kunsagi_rdp_align_nt_clade | *Kunsagivirus* | | | | | **OQ818317** | | | | | | | | | |
| kunsagi_rdp_align_nt_clade | *Kunsagivirus* | 1815 | 1865 | NC_033818.1 | OP589993.1 | NC_034206.1 | 1.54E-03 | NS | NS | NS | NS | NS | 1 | no | |
| kunsagi_rdp_align_nt_clade | *Kunsagivirus* | | | **OQ818317** | | | | | | | | | | | |
| kunsagi_rdp_align_nt_clade | *Kunsagivirus* | 4320 | 4779 | NC_034206.1 | **OQ818317** | Unknown (rodent_china) | 5.16E-03 | NS | NS | NS | NS | NS | 1 | no | |
| kunsagi_rdp_align_nt | *Kunsagivirus* | 5893 | 5949 | NC_038317 | OQ716009 | **Unknown (OQ818317)** | 1.83E-04 | 1.90E-02 | NS | NS | NS | 5.50E-03 | 3 | no | |
| kunsagi_rdp_align_nt | *Kunsagivirus* | | | | | Unknown(NC_033818.1) | | | | | | | | | |
| kunsagi_rdp_align_nt | *Kunsagivirus* | 1814 | 1865 | NC_033818.1 | OP589993.1 | NC_034206.1 | 2.92E-03 | NS | NS | NS | NS | NS | 1 | no | |
| kunsagi_rdp_align_nt | *Kunsagivirus* | | | **OQ818317** | | | | | | | | | | | |
| kunsagi_rdp_align_nt | *Kunsagivirus* | 4893 | 5027 | NC_038317 | OQ716008 | **Unknown (OQ818317)** | 8.00E-03 | NS | NS | NS | NS | 1.99E-02 | 2 | no | |
| kunsagi_rdp_align_nt | *Kunsagivirus* | 4351 | 4670 | NC_034206.1 | NC_038317 | **Unknown (OQ818317)** | NS | NS | 8.82E-03 | 7.05E-03 | 7.58E-03 | NS | 3 | no | |
| kunsagi_rdp_align_nt | *Kunsagivirus* | 3461 | 3734 | NC_033818.1 | Unknown (NC_034206.1) | NC_038317 | 5.24E-03 | 2.60E-02 | 3.30E-02 | NS | 3.44E-02 | NS | 4 | yes | |
| kunsagi_rdp_align_nt | *Kunsagivirus* | | | **OQ818317** | | | | | | | | | | | |
| kunsagi_rdp_align_nt | *Kunsagivirus* | 5966 | 6057 | **OQ818317** | OQ716009 | Unknown (NC_038317) | 1.02E-02 | NS | NS | NS | NS | NS | 1 | no | |
| kunsagi_rdp_align_nt | *Kunsagivirus* | | | | OQ716008 | | | | | | | | | | |
| mischi_rdp_align_nt | *Mischivirus* | 5269 | 5420 | NC_026470.1 | **OQ818316** | MG888045.1 | 6.34E-04 | NS | 3.65E-03 | 8.00E-03 | NS | NS | 3 | no | |
| mischi_rdp_align_nt | *Mischivirus* | 2021 | 2127 | NC_075428.1 | MG888045.1 | **Unknown (OQ818316)** | 1.44E-02 | NS | NS | NS | NS | NS | 1 | no | |
| mischi_rdp_align_nt | *Mischivirus* | 6142 | 6198 | OR951360.1 | NC_043072.1 | **OQ818316** | 3.86E-02 | NS | NS | NS | NS | NS | 1 | no | |
| mischi_rdp_align_nt | *Mischivirus* | 2128 | 2388 | NC_075428.1 | **Unknown (OQ818316)** | NC_043072.1 | 4.12E-02 | NS | NS | NS | NS | NS | 1 | no | |
| sapelo_rdp_align_nt_clade | *Sapelovirus* | 7443 | 269 | PP711943 | Unknown (NC_003987) | **OQ818320** | NS | NS | 1.10E-12 | 4.02E-02 | 2.41E-06 | NS | 3 | no | |
| sapelo_rdp_align_nt_clade | *Sapelovirus* | 268 | 817 | **OQ818321** | Unknown (PP711911) | PP711943 | 1.99E-09 | 1.11E-06 | 7.83E-07 | 2.57E-07 | 2.46E-05 | NS | 5 | yes | Sfig. 5B |
| sapelo_rdp_align_nt_clade | *Sapelovirus* | | | | | NC_033820 | | | | | | | | | |
| sapelo_rdp_align_nt_clade | *Sapelovirus* | | | | | **OQ818320** | | | | | | | | | |
| sapelo_rdp_align_nt_clade | *Sapelovirus* | 3071 | 5027 | NC_033820 | **OQ818320** | PP711943 | 6.34E-09 | NS | 6.77E-06 | 4.66E-10 | 2.87E-11 | 1.87E-14 | 5 | yes | Fig. 4A |
| sapelo_rdp_align_nt_clade | *Sapelovirus* | 101 | 758 | PP711921 | Unknown (PP711911) | NC_033820 | 5.94E-10 | 8.31E-08 | 2.98E-07 | 4.71E-07 | 1.17E-07 | NS | 4 | yes | |
| sapelo_rdp_align_nt_clade | *Sapelovirus* | | | | | **OQ818320** | | | | | | | | | |
| sapelo_rdp_align_nt_clade | *Sapelovirus* | 2991 | 3919 | **OQ818329** | PP711921 | Unknown (PP711943) | 7.65E-08 | NS | NS | 1.39E-02 | 6.74E-07 | NS | 3 | no | |
| sapelo_rdp_align_nt_clade | *Sapelovirus* | | | | **OQ818320** | **Unknown(OQ818321)** | | | | | | | | | |
| sapelo_rdp_align_nt_clade | *Sapelovirus* | | | | PP711943 | Unknown(PP711921) | | | | | | | | | |
| sapelo_rdp_align_nt_clade | *Sapelovirus* | | | **OQ818329** | **OQ818321** | | | | | | | | | | |
| sapelo_rdp_align_nt_clade | *Sapelovirus* | 2236 | 2304 | PP711911 | Unknown (PP711921) | **OQ818320** | 1.34E-04 | 1.34E-03 | NS | 2.27E-03 | 2.30E-02 | NS | 4 | yes | |
| sapelo_rdp_align_nt_clade | *Sapelovirus* | | | | | NC_033820 | | | | | | | | | |
| sapelo_rdp_align_nt_clade | *Sapelovirus* | 6799 | 6863 | PP711921 | **OQ818321** | **Unknown (OQ818320)** | 2.91E-04 | 6.80E-03 | NS | NS | NS | NS | 2 | no | |
| sapelo_rdp_align_nt_clade | *Sapelovirus* | 6476 | 6624 | **OQ818329** | PP711921 | Unknown (NC_003987) | 9.19E-04 | NS | 3.69E-02 | NS | NS | NS | 2 | no | |
| sapelo_rdp_align_nt_clade | *Sapelovirus* | | | PP711943 | | | | | | | | | | | |
| sapelo_rdp_align_nt_clade | *Sapelovirus* | 5119 | 6539 | PP711943 | **Unknown (OQ818329)** | NC_033820 | NS | 1.29E-14 | 5.87E-14 | 5.40E-04 | 4.16E-08 | NS | 4 | yes | |
| sapelo_rdp_align_nt_clade | *Sapelovirus* | | | | | **OQ818320** | | | | | | | | | |
| sapelo_rdp_align_nt_clade | *Sapelovirus* | 3977 | 5118 | PP711943 | **Unknown (OQ818329)** | **OQ818320** | NS | 1.14E-15 | 8.48E-14 | 4.69E-18 | NS | NS | 3 | no | |
| sapelo_rdp_align_nt_clade | *Sapelovirus* | | | **OQ818320** | | | | | | | | | | | |
| sapelo_rdp_align_nt_clade | *Sapelovirus* | 6711 | 7438 | PP711943 | **Unknown (OQ818329)** | **OQ818320** | NS | 8.47E-10 | 2.67E-07 | 3.01E-03 | 1.94E-04 | NS | 4 | yes | |
| sapelo_rdp_align_nt_clade | *Sapelovirus* | 48 | 441 | NC_003987 | **OQ818320** | Unknown (PP711911) | 7.12E-03 | NS | 2.90E-03 | 6.90E-03 | NS | NS | 3 | no | |
| sapelo_rdp_align_nt_clade | *Sapelovirus* | | | | PP711911 | **Unknown(OQ818320)** | | | | | | | | | |
| sapelo_rdp_align_nt_clade | *Sapelovirus* | 2169 | 2281 | **OQ818329** | **Unknown (OQ818321)** | **OQ818320** | 1.25E-02 | NS | NS | NS | NS | NS | 1 | no | |
| sapelo_rdp_align_nt_clade | *Sapelovirus* | | | | | NC_033820 | | | | | | | | | |
| sapelo_rdp_align_nt_clade | *Sapelovirus* | 5647 | 5711 | PP711911 | eonycteris_china | **OQ818320** | 1.54E-02 | NS | NS | NS | NS | NS | 1 | no | |
| sapelo_rdp_align_nt_clade | *Sapelovirus* | 1273 | 1313 | PP711943 | **OQ818321** | NC_003987 | 1.78E-02 | NS | NS | NS | NS | NS | 1 | no | |
| sapelo_rdp_align_nt_clade | *Sapelovirus* | | | | PP711921 | | | | | | | | | | |
| sapelo_rdp_align_nt_clade | *Sapelovirus* | 2408 | 2445 | **OQ818329** | PP711911 | NC_033820 | 3.73E-02 | NS | NS | NS | NS | NS | 1 | no | |
| sapelo_rdp_align_nt_clade | *Sapelovirus* | 6199 | 6458 | **OQ818321** | NC_033820 | PP711911 | 0.04404758 | NS | NS | 1.41E-02 | NS | NS | 2 | no | |
| sapelo_rdp_align_nt_clade | *Sapelovirus* | | | PP711921 | | | | | | | | | | | |
| sapelo_rdp_align_nt_clade | *Sapelovirus* | | | **OQ818321** | | | | | | | | | | | |
| sapelo_rdp_align_nt | *Sapelovirus* | 7443 | 269 | PP711943 | Unknown (NC_003987) | **OQ818320** | NS | NS | 4.52E-12 | 6.59E-09 | 2.04E-05 | NS | 3 | no | |
| sapelo_rdp_align_nt | *Sapelovirus* | 7733 | 847 | **OQ818321** | Unknown (OR951327) | NC_033820 | 5.57E-08 | NS | 4.72E-09 | 7.25E-04 | 6.45E-08 | NS | 4 | yes | |
| sapelo_rdp_align_nt | *Sapelovirus* | | | | Unknown(OR951326) | | | | | | | | | | |
| sapelo_rdp_align_nt | *Sapelovirus* | 3070 | 5027 | NC_033820 | **OQ818320** | PP711943 | 1.66E-08 | NS | 1.77E-05 | 1.22E-09 | 7.52E-11 | 4.88E-14 | 5 | yes | |
| sapelo_rdp_align_nt | *Sapelovirus* | | | | Unknown(OR951327) | **OQ818320** | | | | | | | | | |

| | | | | | | | | | | | | | | | |
|---|---|---|---|---|---|---|---|---|---|---|---|---|---|---|---|
| sapelo_rdp_align_nt | *Sapelovirus* | | | **OQ818321** | Unknown(OR951327) | **OQ818320** | | | | | | | | | |
| sapelo_rdp_align_nt | *Sapelovirus* | 6796 | 6862 | PP711921 | **OQ818321** | **Unknown (OQ818320)** | 1.66E-04 | 9.61E-04 | NS | NS | NS | NS | 2 | no | |
| sapelo_rdp_align_nt | *Sapelovirus* | | | | | Unknown(NC_033820) | | | | | | | | | |
| sapelo_rdp_align_nt | *Sapelovirus* | 2237 | 2303 | PP711911 | Unknown (PP711921) | **OQ818320** | 2.20E-04 | 2.46E-03 | NS | 4.03E-03 | 2.50E-02 | NS | 4 | yes | |
| sapelo_rdp_align_nt | *Sapelovirus* | 4437 | 5660 | PP711921 | Unknown (OR951326) | **OQ818320** | 1.33E-03 | NS | NS | 5.37E-03 | 2.71E-03 | NS | 3 | no | |
| sapelo_rdp_align_nt | *Sapelovirus* | | | **OQ818321** | | PP711943 | | | | | | | | | |
| sapelo_rdp_align_nt | *Sapelovirus* | 6279 | 6429 | NC_003987 | **Unknown (OQ818329)** | PP711921 | 2.32E-03 | NS | NS | NS | NS | NS | 1 | no | |
| sapelo_rdp_align_nt | *Sapelovirus* | 0 | 384 | **OQ818329** | Unknown (NC_003987) | **OQ818320** | 3.39E-03 | 1.78E-02 | 1.36E-02 | 1.38E-04 | NS | NS | 4 | yes | |
| sapelo_rdp_align_nt | *Sapelovirus* | | | PP711911 | | NC_033820 | | | | | | | | | |
| sapelo_rdp_align_nt | *Sapelovirus* | 7170 | 7256 | PP711911 | OR951326 | **OQ818320** | NS | NS | 6.53E-03 | NS | NS | NS | 1 | no | |
| sapelo_rdp_align_nt | *Sapelovirus* | | | | | NC_033820 | | | | | | | | | |
| sapelo_rdp_align_nt | *Sapelovirus* | 3143 | 4199 | **OQ818329** | OR951325 | Unknown (PP711943) | 0.00967749 | NS | NS | NS | NS | NS | 1 | no | |
| sapelo_rdp_align_nt | *Sapelovirus* | | | | PP711943 | Unknown(OR951325) | | | | | | | | | |
| sapelo_rdp_align_nt | *Sapelovirus* | 1773 | 1796 | PP711943 | **OQ818320** | Unknown (OR951332) | 1.40E-02 | NS | NS | NS | NS | NS | 1 | no | |
| sapelo_rdp_align_nt | *Sapelovirus* | 1752 | 1933 | OR951325 | OR951332 | **OQ818320** | 1.44E-02 | NS | 8.25E-03 | 5.35E-03 | 9.49E-03 | NS | 4 | yes | |
| sapelo_rdp_align_nt | *Sapelovirus* | 4182 | 4459 | NC_003987 | OR951325 | Unknown (PP711943) | 0.01191445 | NS | NS | 1.98E-02 | 1.83E-02 | NS | 3 | no | |
| sapelo_rdp_align_nt | *Sapelovirus* | | | | **OQ818320** | Unknown(OR951325) | | | | | | | | | |
| sapelo_rdp_align_nt | *Sapelovirus* | | | | PP711943 | | | | | | | | | | |
| sapelo_rdp_align_nt | *Sapelovirus* | 845 | 1051 | OR951325 | Unknown (PP711921) | PP711943 | 1.79E-02 | NS | NS | NS | NS | NS | 1 | no | |
| sapelo_rdp_align_nt | *Sapelovirus* | 621 | 639 | **OQ818329** | PP711911 | PP711943 | 3.97E-02 | NS | NS | NS | NS | NS | 1 | no | |
| sapelo_rdp_align_nt | *Sapelovirus* | 6465 | 6500 | **OQ818329** | PP711911 | NC_033820 | 4.30E-02 | NS | NS | NS | NS | NS | 1 | no | |
| sapelo_rdp_align_nt | *Sapelovirus* | 2162 | 2547 | **OQ818329** | Unknown (PP711921) | PP711943 | NS | NS | 2.25E-02 | 1.42E-02 | NS | NS | 2 | no | |
| sapo_rdp_align_nt_clade | *Sapovirus* | 1140 | 1451 | e_helvum_cameroon | ON872527 | OQ709197 | 5.70E-03 | NS | 7.73E-03 | 2.58E-02 | NS | NS | 3 | no | |
| sapo_rdp_align_nt_clade | *Sapovirus* | | | **OQ818319** | | | | | | | | | | | |
| sapo_rdp_align_nt_clade | *Sapovirus* | | | **PP766459** | | | | | | | | | | | |
| sapo_rdp_align_nt_clade | *Sapovirus* | | | OP963623 | | | | | | | | | | | |
| sapo_rdp_align_nt_clade | *Sapovirus* | 3225 | 7029 | e_helvum_cameroon | Unknown (r_aegypticus_kenya) | **PP766459** | 4.64E-03 | NS | NS | 3.18E-05 | 3.97E-05 | 3.66E-04 | 4 | yes | |
| sapo_rdp_align_nt_clade | *Sapovirus* | 7749 | 893 | r_aegypticus_kenya | OQ709197 | **PP766459** | 1.29E-02 | NS | 1.21E-02 | 1.97E-02 | 2.03E-04 | NS | 4 | yes | |
| sapo_rdp_align_nt_clade | *Sapovirus* | 196 | 405 | ON872527 | Unknown (OQ709197) | **PP766459** | 3.91E-02 | NS | NS | NS | NS | NS | 1 | no | |
| sapo_rdp_align_nt_clade | *Sapovirus* | 1983 | 2066 | ON872527 | Unknown (OP963622) | **PP766459** | 2.44E-02 | 0.0243913 | NS | NS | NS | NS | 2 | no | |
| sapo_rdp_align_nt_clade | *Sapovirus* | | | | | **OQ818319** | | | | | | | | | |
| sapo_rdp_align_nt_clade | *Sapovirus* | 6645 | 6815 | OP963623 | **PP766459** | OQ709197 | 4.22E-02 | NS | NS | 1.68E-03 | 1.20E-03 | 3.78E-02 | 4 | yes | |
| sapo_rdp_align_nt_clade | *Sapovirus* | | | | **OQ818319** | | | | | | | | | | |
| sapo_rdp_align_nt_clade | *Sapovirus* | 5752 | 6021 | OP963623 | OP963622 | ON872527 | 4.51E-02 | NS | NS | NS | 2.81E-02 | NS | 2 | no | |
| sapo_rdp_align_nt_clade | *Sapovirus* | | | **OQ818319** | | | | | | | | | | | |
| sapo_rdp_align_nt_clade | *Sapovirus* | | | **PP766459** | | | | | | | | | | | |
| sapo_rdp_align_nt | *Sapovirus* | 4883 | 5167 | KX759623 | **PP766459** | PP712001 | NS | NS | 8.36E-04 | NS | NS | 7.22E-03 | 2 | no | |
| sapo_rdp_align_nt | *Sapovirus* | | | | **OQ818319** | | | | | | | | | | |
| sapo_rdp_align_nt | *Sapovirus* | 2870 | 2948 | KX759621 | **PP766459** | OQ709197 | 1.27E-03 | NS | NS | NS | NS | NS | 1 | no | |
| sapo_rdp_align_nt | *Sapovirus* | | | | **OQ818319** | | | | | | | | | | |
| sapo_rdp_align_nt | *Sapovirus* | 6726 | 6787 | OP963622 | **Unknown (PP766459)** | NC_033776 | 1.31E-03 | NS | NS | NS | NS | NS | 1 | no | |
| sapo_rdp_align_nt | *Sapovirus* | | | | **Unknown(OQ818319)** | KX759622 | | | | | | | | | |
| sapo_rdp_align_nt | *Sapovirus* | 925 | 1501 | KX759619 | Unknown (PP712001) | **PP766459** | 1.88E-03 | NS | 3.84E-02 | NS | NS | NS | 2 | no | |
| sapo_rdp_align_nt | *Sapovirus* | | | KX759623 | | **OQ818319** | | | | | | | | | |
| sapo_rdp_align_nt | *Sapovirus* | 1111 | 1399 | KX759623 | ON872527 | OQ709197 | 3.11E-03 | NS | NS | 3.16E-02 | 0.03500671 | NS | 3 | no | |
| sapo_rdp_align_nt | *Sapovirus* | | | **OQ818319** | | | | | | | | | | | |
| sapo_rdp_align_nt | *Sapovirus* | | | **PP766459** | | | | | | | | | | | |
| sapo_rdp_align_nt | *Sapovirus* | 1988 | 2076 | OQ709197 | Unknown (PP712004) | **PP766459** | 1.40E-02 | NS | NS | NS | NS | NS | 1 | no | |
| sapo_rdp_align_nt | *Sapovirus* | | | | | **OQ818319** | | | | | | | | | |
| sapo_rdp_align_nt | *Sapovirus* | 4363 | 4565 | KX759619 | Unknown (PP712001) | **PP766459** | 2.86E-02 | NS | NS | NS | NS | NS | 1 | no | |
| sapo_rdp_align_nt | *Sapovirus* | | | | **Unknown(OQ818319)** | PP712001 | | | | | | | | | |
| sapo_rdp_align_nt | *Sapovirus* | | | | **Unknown(PP766459)** | | | | | | | | | | |
| tescho_rdp_align_nt_clade | *Teschovirus* | 6968 | 7180 | r_aegypticus_africa | **OQ818323** | **Unknown (OQ818324)** | 1.04E-20 | 3.09E-07 | 4.54E-21 | 9.73E-11 | 6.32E-10 | 8.05E-09 | 6 | yes | Sfig. 5D |
| tescho_rdp_align_nt_clade | *Teschovirus* | | | | **OQ818324** | **Unknown(OQ818323)** | | | | | | | | | |
| tescho_rdp_align_nt_clade | *Teschovirus* | 28 | 915 | **OQ818323** | **OQ818324** | r_aegypticus_africa | 4.70E-06 | NS | 5.47E-04 | 1.19E-08 | 1.76E-06 | 1.66E-04 | 5 | yes | Sfig. 5C |
| tescho_rdp_align_nt_clade | *Teschovirus* | 1128 | 1887 | rousettus_china | Unknown (OR951324) | **OQ818318** | 6.19E-03 | NS | 1.06E-02 | 3.34E-02 | 0.04247208 | NS | 4 | yes | |
| tescho_rdp_align_nt_clade | *Teschovirus* | 3299 | 6944 | r_aegypticus_africa | **OQ818324** | **Unknown (OQ818323)** | 2.12E-02 | NS | NS | 1.96E-04 | NS | 1.63E-04 | 3 | no | |
| tescho_rdp_align_nt_clade | *Teschovirus* | | | | **OQ818323** | **Unknown(OQ818324)** | | | | | | | | | |
| tescho_rdp_align_nt_clade | *Teschovirus* | 6736 | 6849 | **OQ818318** | rousettus_china | LC386158 | NS | NS | 4.14E-02 | 0.02157041 | NS | NS | 2 | no | |
| tescho_rdp_align_nt | *Teschovirus* | 5039 | 913 | PP711934 | **OQ818324** | **OQ818323** | 2.33E-15 | NS | 6.56E-06 | 2.02E-04 | 5.00E-10 | 2.66E-14 | 5 | yes | Fig. 4B |
| tescho_rdp_align_nt | *Teschovirus* | 3231 | 5072 | **OQ818323** | Unknown (OR951335) | PP711934 | 1.63E-07 | NS | 9.02E-08 | 2.11E-02 | 2.88E-10 | NS | 4 | yes | |
| tescho_rdp_align_nt | *Teschovirus* | 6692 | 6832 | PP711948 | **OQ818318** | LC386158 | 1.56E-04 | NS | 9.86E-03 | NS | NS | 9.78E-05 | 3 | no | |
| tescho_rdp_align_nt | *Teschovirus* | 539 | 749 | OR951335 | **Unknown (OQ818324)** | **OQ818323** | 2.20E-03 | NS | NS | 3.81E-03 | 2.02E-02 | NS | 3 | no | |
| tescho_rdp_align_nt | *Teschovirus* | 6307 | 54 | OR951335 | Unknown (OR951333) | **OQ818323** | 2.24E-03 | NS | NS | 4.51E-05 | 1.81E-02 | NS | 3 | no | |
| tescho_rdp_align_nt | *Teschovirus* | 3981 | 4236 | OR951324 | **OQ818323** | OR951334 | 4.02E-03 | NS | NS | NS | NS | NS | 1 | no | |
| tescho_rdp_align_nt | *Teschovirus* | | | **OQ818318** | | | | | | | | | | | |
| tescho_rdp_align_nt | *Teschovirus* | 3534 | 3708 | **OQ818318** | PP711948 | OR951334 | NS | NS | 1.87E-02 | NS | NS | NS | 1 | no | |
| tescho_rdp_align_nt | *Teschovirus* | 6284 | 6306 | OR951335 | **OQ818324** | OR951333 | NS | NS | 2.55E-02 | NS | NS | NS | 1 | no | |
| tescho_rdp_align_nt | *Teschovirus* | 74 | 354 | OR951335 | **OQ818324** | Unknown (OR951333) | NS | NS | 2.24E-02 | NS | 2.17E-02 | NS | 2 | no | |
| tescho_rdp_align_nt | *Teschovirus* | 1965 | 2029 | **OQ818323** | PP711934 | **Unknown (OQ818318)** | 2.56E-02 | NS | NS | NS | NS | NS | 1 | no | |
| tescho_rdp_align_nt | *Teschovirus* | 1482 | 1660 | OR951333 | OR951324 | **OQ818324** | 3.02E-02 | NS | 2.87E-02 | 0.04413962 | NS | NS | 3 | no | |
| tescho_rdp_align_nt | *Teschovirus* | | | OR951334 | | | | | | | | | | | |
| tescho_rdp_align_nt | *Teschovirus* | 1237 | 1321 | OR951335 | **OQ818323** | **OQ818318** | 3.22E-02 | NS | NS | NS | NS | NS | 1 | no | |
